# Identification and proteomic profiling of CD90^+^ small EVs using a refined immunocapture separation approach targeting stromal-derived EV subpopulations in synovial fluid of arthritis patients

**DOI:** 10.1101/2025.03.12.640750

**Authors:** Stefanie Kurth, Edveena Hanser, Simone Häner-Massimi, Stavros Giaglis, Florian Geier, Dominik Burri, Katarzyna Buczak, Ute Heider, Stefan Wild, André N. Tiaden, Diego Kyburz

## Abstract

Extracellular vesicles (EVs) have emerged as essential drivers in disease progression and promising biomarkers across various disease conditions, including autoimmune diseases such as rheumatoid arthritis (RA). RA is characterized by chronic inflammation driven by patient-specific cellular sub-phenotypes within the synovium. Since EVs reflect their cellular origin and are detectable in biofluids like synovial fluid, they offer a promising alternative for novel biomarker discovery or diagnostics in arthritis, potentially reducing the need for invasive tissue biopsies. However, the inherent complexity, viscosity and extracellular matrix (ECM) composition of synovial fluid pose significant challenges for efficient EV isolation. Moreover, most previous studies have focused on heterogenous bulk EV populations, neglecting the diverse EV subsets present in patient biofluids.

To address these challenges, we established a novel immunocapture-based separation method, that selectively targets and isolates cell type-specific EV subpopulation from the heterogenous EV pool present in synovial fluid of arthritis patients. As a proof-of-concept approach, we targeted a stromal-derived EV population by leveraging the fibroblast-associated surface marker CD90/THY1, representing a subset of synovial fibroblasts implicated as important drivers in synovitis and chronic inflammation in RA. Western Blot and data independent acquisition (DIA) liquid chromatography coupled to tandem mass spectrometry (LC-MS/MS) proteomic analysis of immunocaptured CD90^+^ EVs isolated from cultured primary synovial fibroblasts and synovial fluid of arthritis patients, confirmed the successful separation of stromal vs a myeloid/lymphoid EV populations. The in-depth proteomic profiling of synovial fluid EVs offers unprecedented insights into EV heterogeneity in arthritis. This analysis enables the classification of cell-type associated EV subsets and the identification of novel EV markers for future immunocapture-based EV separation strategies. Our refined fit-for-purpose EV separation approach, coupled to in depth proteomic profiling, provides a powerful tool to identify disease-relevant EV subpopulations and novel biomarker candidates in synovial fluid, facilitating advances in arthritis diagnostics and personalized medicine

## Introduction

### EV function and use as biomarker

Extracellular vesicles (EV) represent a heterogenous ensemble of nanosized membrane-bound particles ubiquitously secreted across diverse cell types^1^. Research in the last decade established EVs as instrumental entities in intercellular communication which facilitate component exchange between cells in their vicinity but also convey functional cargo to distant tissues and organs^2,3^. EVs are pivotal in mediating a broad spectrum of biological processes including differentiation, proliferation and regeneration or supporting cellular and tissue homeostasis by quickly adapting to environmental changes^4^. Recently, immunomodulatory functions were attributed to EVs derived from the immune system, including antigen presentation, activation of immune cells, regulation of inflammation or immune surveillance, thereby underscoring their significance in health and disease^5–8^. As such, EVs are implicated in the aetiology of numerous pathological conditions involving the immune system, particularly in cancer and in complex diseases originating from dysfunctional immune regulation^9,10^. In chronic inflammatory arthritis, such as Rheumatoid arthritis (RA), EVs were shown to have an instrumental function in disease progression^11,12^. EVs generated from chronically inflamed joint tissue contribute to sustained inflammation by downregulating anti-inflammatory genes via transferred miRNAs, associating with citrullinated proteins generating active immune-complexes, promoting angiogenesis or inhibiting apoptosis of the stromal compartment^13–15^. Consequentially, EVs have rapidly emerged as novel therapeutic targets and promising biomarker candidates, offering novel perspectives to identify disease-relevant mechanisms and monitor efficacy of selected therapies^16–18^.

The utility of EVs as potential biomarkers is primarily attributed to their unique molecular composition of membrane-bound proteins and intravesicular biomolecules, which both meticulously reflect the cellular identity and the physiological status of their tissue of origin^19^. Thus, EVs carry a diverse array of bioactive molecules integral to numerous physiological processes, including nucleic acids (miRNA, mRNA, DNA), proteins, signalling components, transcription factors and metabolites^20^. in addition, the secreted vesicles maintain the same membrane orientation as their parental cell, often incorporating cell type-specific surface proteins and lipids^21^. In the context of RA, EVs are anticipated to carry molecular signatures that provide a unique perspective on the cellular dynamics in the inflamed tissue^22^. Thus, analysing the functional surfaceome of the EV membrane and the encapsulated cargo in the EV lumen derived from RA patients offer a window into the current pathophysiological processes, providing a real-time snapshot of disease activity and progression, with significant implications for diagnosis, prognosis, and monitoring of therapeutic responses. Furthermore, the omni-presence of EVs and their exceptional stability in biological fluids renders them valuable for non-invasive biomarker discovery ^23^, especially in a disease condition like arthritis where direct tissue sampling from the joint is challenging.

### Distinct synovial cell populations drive inflammation in arthritis

Arthropathies like Rheumatoid Arthritis (RA) and Osteoarthritis (OA) represent two frequent manifestations of autoimmune and degenerative disorders in the musculoskeletal disease area, which commonly impact the joints and surrounding tissues, resulting in chronic degeneration of cartilage and bone, pain, and in the case of RA, severe systemic comorbidities ^24–26^. The etiology of RA and OA are multifaceted, including genetic predisposition, environmental influences, individual life style and aging, still the and underlying pathogenesis differs significantly. In RA, a series of yet undefined processes initially disrupt the immune tolerance, followed by an undefined latency period before clinical manifestations can be diagnosed. In contrast, physical overloading, excessive strain and trauma stand at the onset of OA. Both diseases, though, manifest as low- (OA) or high-grade (RA) joint-centric inflammation^27–30^, with synovitis, the chronic inflammation of the synovial membrane being a pathological hallmark in both conditions. Synovitis is driven and sustained via interactions between distinct subsets of synovial cells, mainly stromal cells and tissue-resident macrophages, and cells of the periarticular tissue, like chondrocytes, adipocytes or osteoblasts^31–33^. The aberrant activation of these tissue-resident cells manifests as uncontrolled inflammatory circuits which in turn drive the recruitment, infiltration and activation of circulating immune cells into the tissue. This cascade of paracrine and autocrine activation due to the exuberate immune response perpetuates the chronicity of synovitis resulting in dysfunctional joint homeostasis, increased angiogenesis, and ultimately progressive degeneration of articular cartilage and bone^34^.

Recent advancements in molecular profiling of synovial membrane tissue by multimodal single cell omics approaches, particularly single-cell RNA sequencing (scRNA-Seq) studies in the last 5 years^35^, revealed distinct histopathological phenotypes that correlate with disease activity and treatment outcomes^36–38^. The holistic analysis of the single cell gene expression profiles in RA synovial tissue led to the identification of novel functional subsets of synovium-located stromal, myeloid and lymphoid cells - these cellular clusters were referred to as “cell-type abundance phenotypes” (CTAPs), each characterized by selectively enriched cell states^39,40^. Synovial fibroblasts are key cells in the synovium driving and perpetuating synovitis^41^, and CTAPs dominated by stromal cells are exhibiting distinct functional fibroblast subsets each characterized by an ensemble of cellular and molecular markers^42,43^. Hence, a specific subset identified by CD90^-^PDPN^+^FAPalpha^+^ expression is implicated in joint destruction through the secretion of catabolic extracellular matrix (ECM) proteases, while a CD90^+^PDPN^+^FAPalpha^+^- subset is contributing to the perpetuation of inflammatory conditions by releasing proinflammatory cytokines^44–47^.

### EVs in synovial fluid represent physiological/pathological conditions of inflamed synovial tissue

Although these newly characterized synovial cells hold promises as indicators of disease activity^48,49^, their potential clinical utility currently depends on invasive surgical intervention or needle biopsies to obtain synovial tissue from inflamed joint ^29,50–52^. These procedures are not only costly and require specialized clinical expertise, but also impose significant discomfort on patients. An emerging, less invasive alternative in disease monitoring involves the analysis of robust biomarkers from liquid biopsies, such as synovial fluid from the joint cavity^53,54^. Synovial fluid, a protein-rich exudate produced by the synovial intima, is reflecting the biochemical milieu of the inflamed joint, thus serving as a direct indicator of its pathological state^55–57^. Studies in RA patients have demonstrated that EVs are actively secreted into the synovial fluid by both tissue-resident cells as well as infiltrating immune cells located within the periarticular tissue^11,58,59^. The detection of specific EV subsets in synovial fluid, indicative of the presence of synovitis-associated cellular subsets active in the diseased tissue, provides a new opportunity for a less invasive procedure to identify and follow disease activity.

### EV heterogeneity represents a major challenge for their use as biomarkers

The physical and biochemical nature of synovial fluid as joint lubricant rich of ECM proteins and serum components pose technical challenges for EV isolations, resulting in inadequate yield and co-isolation of a considerable amount of adsorbed non-EV proteins ^60–63^. In addition, EVs derived from patient synovial fluid represent a complex heterogenous mixture of different vesicle populations, varying in size, density, biogenesis, lipid biochemistry and molecular content of encapsulated cargo^64^. This heterogeneity poses a significant challenge for their application as biomarkers in RA or OA. This complexity is further amplified by the disease’s variable clinical manifestations and the dynamic nature of its inflammatory processes. Consequently, the isolation and characterization of specific EV subpopulations pertinent to RA or OA pathogenesis and progression become crucial for their effective use as biomarkers. Addressing these challenges require the development of novel technical approaches or the refinement of present EV isolation and characterization techniques. Existing methodologies, primarily basing on bulk processing, often lack specificity and are susceptible to contaminations with non-EV proteins, and are prone to inaccurate estimations or loss of relevant EV species. Addressing these limitations necessitates the development of targeted approaches, such as immunocapture-based separation, which leverage disease-specific surface marker and provide the ability to selectively isolate EV subpopulations from heterogeneous mixtures. Such advancements are aiming to improve the purity and precision for EV populations, which are critical for ensuring the reliability and clinical utility of EV-based biomarkers in RA and OA.

To overcome the challenges associated with EV heterogeneity and synovial fluid complexity, we introduce a novel, fit-for-purpose EV separation method that integrates nanosized magnetic-bead immunocapture with size exclusion chromatography (SEC) and ultrafiltration (UF). This refined approach enables the selective isolation of synovitis-relevant EV subpopulations from synovial fluid of RA and OA patients, paving the way for improved biomarker discovery and disease monitoring.

## Results

### 1.) Inter-patient variability of marker expressed on synovial fluid derived EVs

The identification of disease-specific EV populations in liquid biopsies has gained significant attention due to their potential roles in disease-mechanisms and as novel biomarkers for chronic inflammatory diseases like RA. To explore and differentiate distinct EV populations present in synovial fluid from arthritis patients, we selected a panel of relevant cell surface markers that are reported to represent abundant pathological cell populations in RA synovial tissue, focusing on the stromal and myeloid compartments^40^. Marker selection was guided by cellular specificity (fibroblast vs. macrophage origin), expression intensity, and the location of the protein in the plasma membrane of the cell, thereby providing the essential criteria for the planned separation of EVs into stromal and myeloid subpopulations such as the accessibility of the marker candidate on the EV surface for antibody binding. For stromal EVs, we targeted THY1 (CD90) and Podoplanin (PDPN), both well-established markers exemplifying stromal phenotypes in RA^47^. Of note, CD90 was previously identified through mass-spectrometry in synovial fluid EVs^65^. For macrophage-related EVs, we selected CD68, CD163 and CD48 representing different macrophages subpopulations within the synovial membrane of RA patients^66,67^.

To assess marker expression, we initially applied a bulk EV isolation protocol using ultracentrifugation (UC), a straightforward and less material-intensive pre-screening method (Figure 1a). Synovial fluid samples from patients with various types of arthritis were analyzed (Figure 1b). As anticipated, the heterogeneity among arthritis patients was evident in the tested marker profiles of the synovial fluid-derived EVs. CD90, a marker linked to pro-inflammatory fibroblast properties in RA synovium, exhibited significant inter-patient variability. In contrast, PDPN, a pan-fibroblast marker, was consistently detected on EVs across all patients. Similarly, macrophage-related markers CD68 and CD48 demonstrated inter-patient variability, whereas CD163, indicative for tissue-resident macrophages, was consistently present in all synovial fluid samples (Figure 1b).

**Figure 1:**
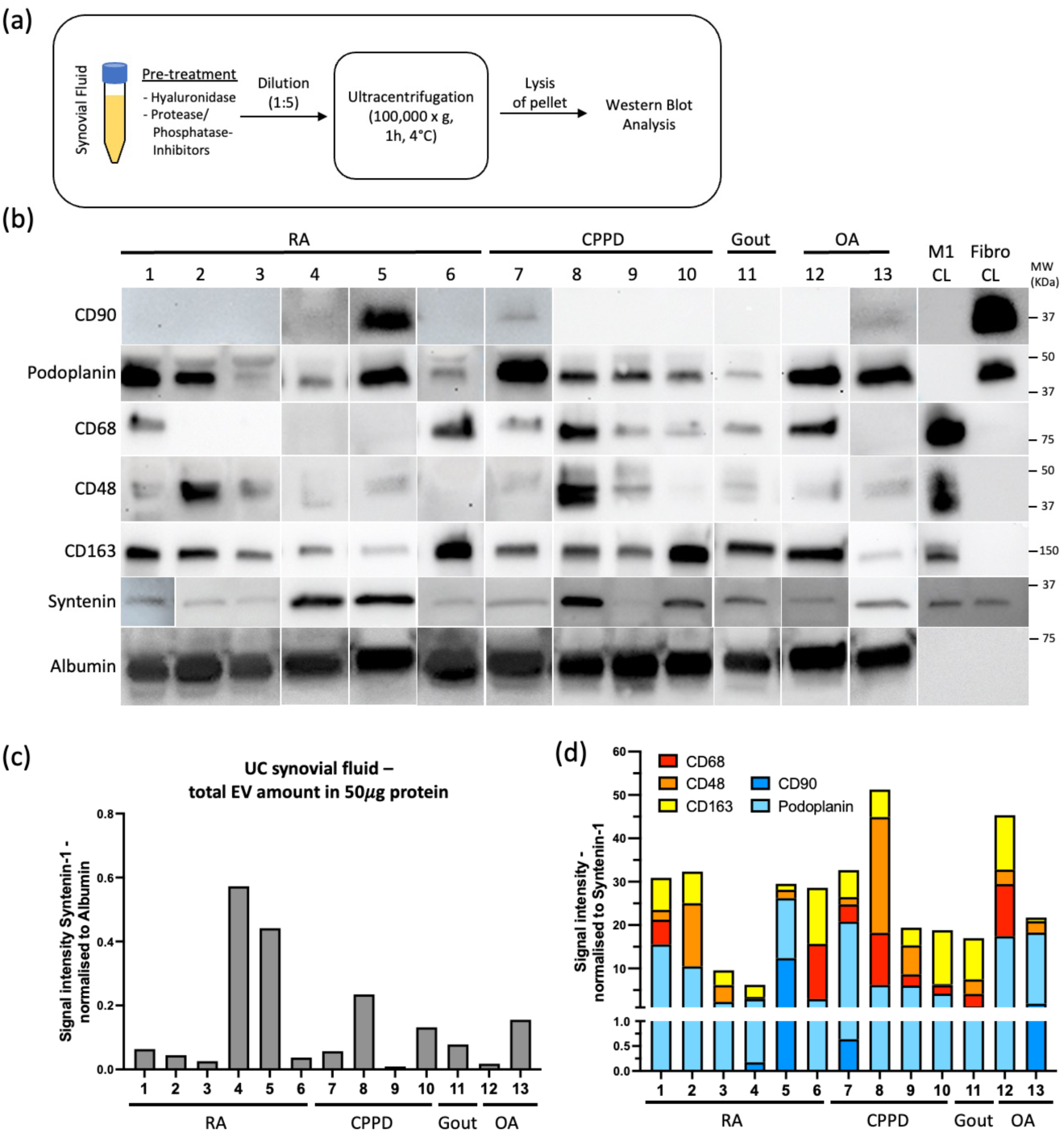
Inter-patient variation of total EV protein and EV surface marker expression in synovial fluid of different arthritis patients. (a) Workflow depicting ultracentrifugation (UC)-generated EV pellets (100’000 x g, 1h, 4°C) from 1ml cell-free synovial fluid of 13 different arthritis patients (6x RA, 4x Calcium pyrophosphate deposition (CPPD) arthritis, 1x gout, 2xOA), diluted 1 to 5 in PBS, treated with hyaluronidase and protease/phosphatase inhibitor, lysed in RIPA buffer and prepared at equal amounts for Western Blot analysis. (b) 50 μg of total protein loaded on SDS-PAGE gels for Western Blot analysis using specific antibodies against selected surface markers (CD90, PDPN, CD68, CD48, CD163, Syntenin-1, human serum albumin). Cell lysates (CL) from cultured macrophages (M1) and synovial fibroblasts (Fibro) (both 2.5 μg) served as controls. (c) Evaluation of EV amount in the UC-isolated samples by normalizing signal intensities of Syntenin-1 (general EV marker, correlates to EV amount) to serum albumin in Western Blot. (d) Distribution of surface marker expression on EVs between different arthritis patient samples by assessing the specific EV marker signal intensity normalized to corresponding Syntenin-1 intensity in Western Blot.

This patient heterogeneity was also reflected in the physical characteristics of synovial fluid, which showed substantial differences in viscosity, color and total protein content present in UC-isolated EV pellets generated from a defined volume (Supplementary Figure S1). These intrinsic differences necessitated normalization approaches to accurately compare EV marker abundance between patient samples. Since analyzing the same quantity of proteins does not necessarily equate to analyzing an equivalent EV population, we employed Syntenin-1 as protein normalizer to quantify Western Blot signal intensity. Syntenin-1 has been considered as universal EV-specific marker due to its consistent abundance across various cellular sources, species, and biofluids, making it an appropriate standard for EV quantification^68^. Accordingly, Western Blot signal intensities were normalized using Syntenin-1 to estimate the ratio of EV-derived protein to total protein in UC synovial fluid isolates (Figure 1c) and to determine the proportion of specific EV markers relative to the total EV protein (Figure 1d). Additionally, serum albumin, considered as a non-EV-derived protein, was used to evaluate the proportion of co-enriched plasma proteins in the synovial fluid samples. While albumin levels were consistent across patient samples, the variability in synovial fluid-derived EV isolates was evident in the differences in Syntenin-1 levels (Figure 1c). Normalizing EV marker expression (PDPN, CD90, CD68, CD48, CD163) to Syntenin-1 provided a more precise evaluation of EV-specific marker abundance, highlighting substantial inter-patient variability (Figure 1d, Supplementary Figure S1). Among the markers analyzed, CD90 showed the least consistent expression on EVs across patients. This variability underscores the complexity of arthritis-related EV populations and highlights the importance of precise normalization and characterization strategies for identifying disease-relevant EV subsets.

### 2.) Size exclusion chromatography coupled with ultrafiltration-based (SEC-UF) Isolation and characterization of RA and OA synovial fibroblast-derived CD90^+^- and PDPN^+^ EVs

Due The significant role of CD90^+^ fibroblasts in RA synovitis and the reported presence of CD90 in synovial fluid EVs, as confirmed in our crude EV UC isolates from arthritis patient SF samples (Figure 1b, d), identified CD90 as a promising synovial EV marker for separating an arthritis-associated EV subpopulation. CD90 (THY-1) encodes a GPI-anchored protein located in the outer leaflet of lipid rafts on stromal cell membranes. Additionally, PDPN, a type I transmembrane sialomucin-like glycoprotein, was selected as pan-synovial EV marker due to its consistent presence in SF-derived EVs. Both proteins are heavily glycosylated and contribute to defining cellular phenotypes through protein-protein interactions, particularly in cancer and immune responses^69,70^.

To assess these markers in arthritis-associated EVs, we first analyzed CD90 and PDPN expression in *ex vivo* cultivated synovial fibroblasts isolated from tissue biopsies of RA and OA patient. Subsequently, we confirmed their presence on small EVs isolated from synovial fibroblasts culture supernatants (SN) using a size exclusion chromatography (SEC) and ultrafiltration (UF)- based protocol (Figure 2a). Transmission electron microscopy (TEM) analysis demonstrated that SEC-UF isolated EVs from RA and OA fibroblast culture SN were intact, round-shaped small vesicles in the size range between 30-130 nm (Figure 2b).

**Figure 2:**
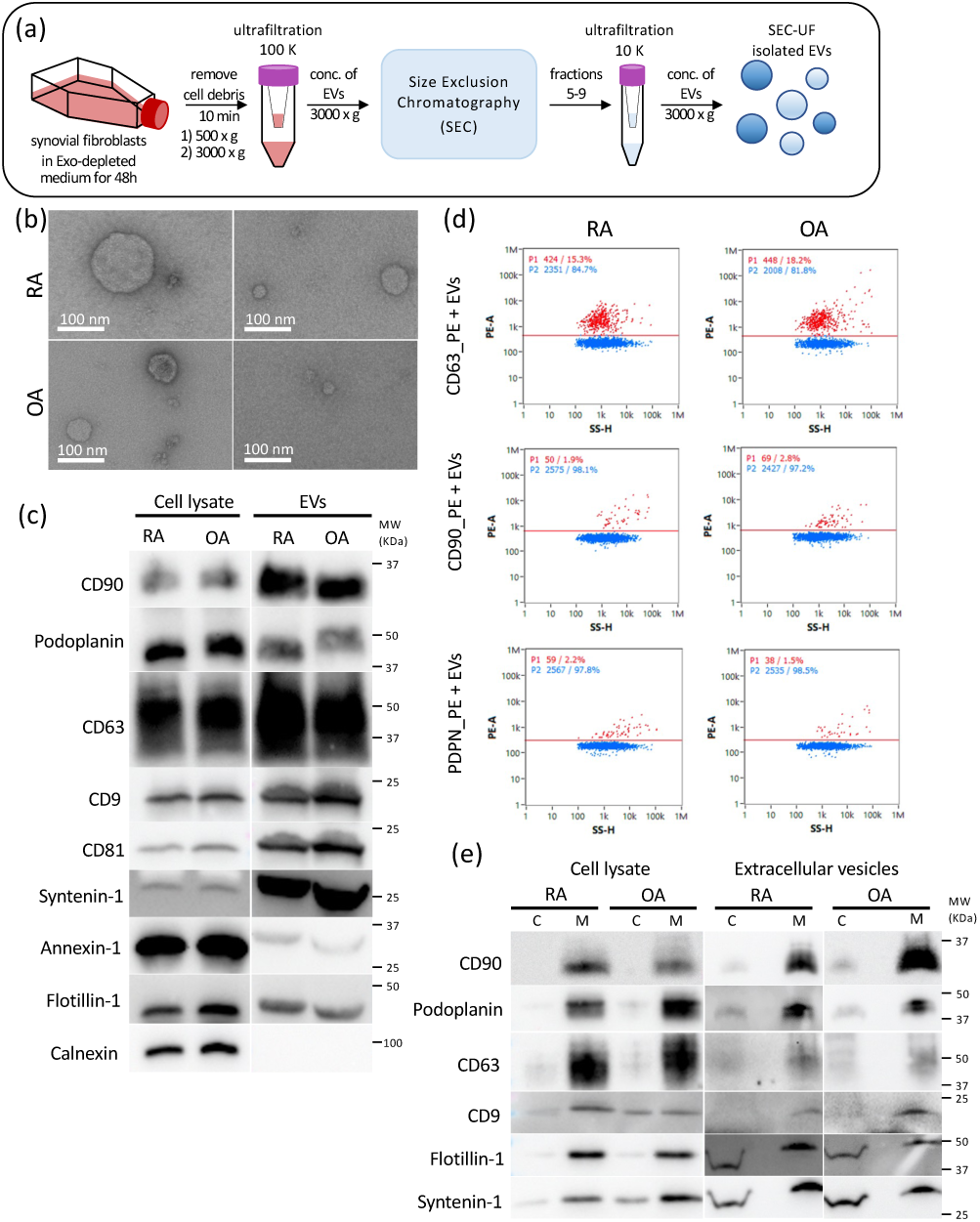
Synovial fibroblast cell culture medium derived EVs isolated using a size exclusion chromatography coupled to ultrafiltration (SEC-UF) protocol contain CD90, Podoplanin and typical small EV markers. (a) Graphical illustration of SEC-UF EV-isolation procedure. The supernatant of patient-derived synovial fibroblasts cultured for 48 h in Exo-depleted medium was centrifugated twice (500 x g 10 min, 3000 x g 10 min) to remove dead cells and cell debris. Cleared supernatant was ultrafiltered using a 100kDa cut-off Amicon column, before applying to the size exclusion chromatography (SEC) using an iZONE qEV1-35nm column. SEC fosters the division of particles regarding their size allowing to sample small EVs by collecting defined fractions. Fractions 5-9 were pooled and ultrafiltered/concentrated using a 10kDA cut-off Amicon column. (b) Representing TEM images of isolated small EVs derived from synovial fibroblasts cell culture (OA vs. RA) ranging from 30-130 nm. (c) Characterization of synovial fibroblast derived SEC-UF isolated EVs via Western Blot analysis. Loading of equal amount of lysed EV protein (4 μg) and corresponding cell lysate (2.5 μg) on a 4-15% SDS gradient gel and analyzed by Western Blot using specific antibodies detecting typical small EV markers (CD63, CD9, CD81, Syntenin-1, Annexin-1, Flotillin-1) and fibroblast markers (CD90, Podoplanin). Calnexin staining was added as EV negative control. (d) NanoFCM measurements of CD63, CD90 and Podoplanin antibody staining of synovial fibroblast cell culture derived EVs. Controls in the supplement (Supplement Figure 2). (e) Western Blot confirming membrane-associated location of CD90 and PDPN in lysates of fibroblasts and fibroblasts-derived EVs using MemPER-treatment separating the cytosolic (C) vs. membrane (M) protein fraction.

Both, RA and OA synovial fibroblasts, express CD90 and PDPN, consistent with previous reports from synovial fibroblast studies and recent synovial tissue profiling data^44,47^. Western Blot analysis confirmed the presence of CD90 and PDPN on EVs isolated from these fibroblasts, alongside typical EV markers, including CD63, CD9, CD81, Syntenin-1, Annexin-1 and Flotillin-1. Calnexin, a cytoplasmic protein predominantly localized in the endoplasmic reticulum, was used as negative control. A strong calnexin signal was observed in the cell lysates but was absent in the isolated EVs, verifying the neatness of the SEC-UF isolation process and the purity of our isolated EVs (Figure 2c).

NanoFCM analysis, flow-cytometry on the nano-metric scale, further validated the presence of CD90 and PDPN on SEC-UF-isolated EVs and their size distribution measured by TEM (30-130 nm) (Figure 2d). NanoFCM also measured EV concentration at 1-2 x 10^10^ EVs/ml, which correlated with similar values obtained through classical NTA measurements (Supplementary Figure S2).

An immunocapture-based separation of intact EV subpopulations relies on a membrane-associated EV surface marker with outward orientation of the antigen region accessible for antibody-binding. The direct interaction of antibodies and membrane-localized CD90 and PDPN was confirmed by positive single-surface staining of EVs using NanoFCM under non-fix/non-permeabilization conditions (Figure 2d) To further verify the membrane localization of CD90 and PDPN, we employed a defined protein lysis protocol to separate the cytosolic and membrane protein fractions of SEC-UF isolated EVs. The results confirmed that the majority of CD90 and PDPN were localized within the EV membrane fraction, consistent with their membrane localization in cell lysates (Figure 2e).

In summary, the SEC-UF isolation and characterization approach confirmed the presence of CD90 and PDPN on the surface of small EVs derived from *ex vivo* cultured synovial fibroblasts. These findings highlight CD90 and PDPN as promising target markers for immunocapture-based techniques to isolate and separate CD90^+^ and PDPN^+^ EV subpopulations. This refined isolation and characterization strategy holds significant potential for studying disease-relevant EV subsets in RA and OA.

### 3.) Separation of CD90^+^ and PDPN^+^ EV subpopulations based on a SEC-UF coupled immunocapture approach with prototype nano-sized magnetic beads

The lack of specific methods for isolating and characterizing distinct EV subpopulations from complex biofluids poses a major challenge in understanding the roles of disease-relevant EV populations. Most studies investigating EVs are based on conventional bulk EV isolation techniques, which recover a heterogenous population of vesicles, and can therefore obscure insights into discrete subpopulations originating from specific disease relevant cell types. To address this limitation, we developed a refined EV separation strategy combining SEC-UF enrichment with immunocapture using prototype nanosized magnetic beads, to selectively target and isolate arthritis-associated EV subpopulation from synovial fluids for subsequent proteomic profiling of disease-relevant cargo.

For our proof-of-principle experiments, we used CD90 and PDPN as initial candidate markers. These markers were confirmed to be present on synovial fibroblast-derived EVs in vitro, and accessible for antibody binding. Using conditioned medium from cultured CD90^+^/PDPN^+^ synovial fibroblast, we employed prototype anti-CD90 or anti-PDPN nanosized magnetic beads (prototype MicroBeads provided by Miltenyi Biotec) to separate either CD90^+^ or PDPN^+^ EVs from SEC-UF enriched EV population, using a modified μMACS separation protocol (Figure 3a).

**Figure 3:**
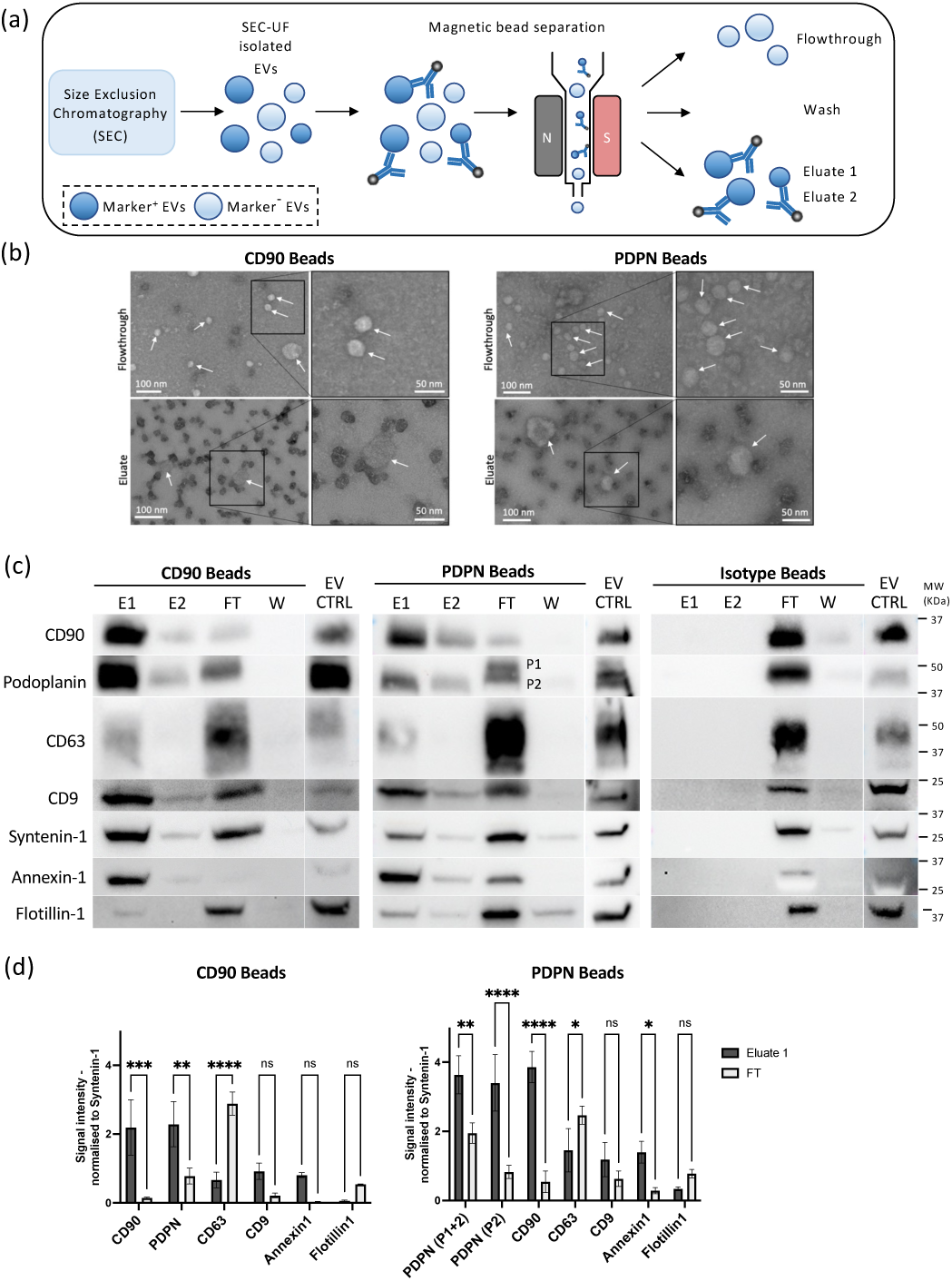
Immunocapture of CD90^+^ and PDPN^+^ EV population. (a) Graphical illustration of immunocapture EV separation approach using prototype nano-sized magnetic beads (EV MicroBeads, Miltenyi). SEC-UF isolated EVs derived from supernatant of in vitro cultured synovial fibroblasts were incubated either with magnetic prototype MicroBeads coupled to anti-CD90 or anti-Podoplanin (PDPN) antibodies or an antibody isotype-coupled bead control (all provided by Miltenyi). Antibody-bound EVs (Marker^+^) are retained by the magnetic column, while unbound EVs (Marker^-^) are collected in the flowthrough (FT). A washing step (W) removed the remaining unbound/loosely associated EVs and protein aggregates. The removal of the magnet, released the bead-bound Marker^+^ EVs, which were collected as Eluate (E1). The elution step was repeated (E2) to evaluate the efficacy of first elution step. (b) TEM measurement of eluate and flowthrough after CD90 and PDPN MicroBead separation. Arrows direct to EVs. (c) Western Blot analysis of CD90 and PDPN MicroBead separations and Isotype control. Detection of CD90, PDPN and typical EV marker (CD63, CD9, Syntenin-1, Annexin-1, Flotillin-1). EV Input (SEC-isolated EVs before separation) was loaded as control. (d) Quantification of CD90 and Podoplanin MicroBead separation from Western Blot results. Quantification of both PDPN populations (P1+P2) and only P2. Normalized to Syntenin-1 amount. 2way ANOVA, multiple comparisons, uncorrected Fishers’s LSD. (* p<0.05,** p<0.005, *** p<0.0005, **** p<0.0001)

#### Magnetic Immunocapture and EV Enrichment

The 50 nm magnetic beads, incubated with 1–2 × 10^10^ EVs for 1 hour at room temperature, demonstrated efficient binding to EV surface markers while avoiding issues such as precipitation and nonspecific protein contamination commonly associated with larger beads (>1 μm). During magnetic separation, bead-bound CD90+ or PDPN+ EVs were retained in the μ-column, while unbound EVs, free proteins, and aggregates passed through and were collected in the flowthrough. Successive washing steps removed loosely bound EVs and nonspecific proteins, while elution yielded enriched CD90^+^ or PDPN^+^ EV populations in two consecutive eluate fractions (eluates 1 and 2); the second eluate (eluate 2) served as a control for the efficiency of the first elution step.

TEM analysis of the eluates confirmed the presence of intact, round-shaped vesicles predominantly bound to magnetic beads, while the flowthrough contained unbound EVs with minimal number free beads not associated with EVs (Figure 3b). Western Blot results revealed a prominent anti-CD90 signal in the eluate, contrasting with a significant weaker signal in the flowthrough confirming the successful enrichment of CD90^+^ EVs (Figure 3c, left panel). Syntenin-1 was present in both fractions facilitating the quantitative normalization of the specific protein signals (Figure 3d, left panel). The Western Blot quantification highlighted distinct characteristics of the CD90^+^ vs. CD90^-^ populations. CD90^+^ EVs were enriched for PDPN, Annexin-1 and CD9 in CD90^+^ EVs, whereas CD63 and Flotillin-1 were more abundant in the CD90^-^ population. In contrast, the isotype-bead control displayed no detectable signal in the eluate but was exclusively present in the flowthrough (Figure 3c, right panel), confirming the specificity of the bead-coupled anti-CD90 antibody. The PDPN-based separation yielded similar results, identifying a PDPN^+^ EV subpopulation co-expressing CD90 and enriching Annexin-1 and CD9, while CD63 and Flotillin-1 were again elevated in the PDPN^-^ population (Figure 3c, middle panel). However, PDPN enrichment in the eluate fraction was less clear. Although there was an enrichment of PDPN in the eluate, residual PDPN signals at a considerable intensity was detectable in the flowthrough indicating potential variation in PDPN protein accessibility or modification (Figure 3d, right panel). Notably, the PDPN protein in the flowthrough exhibited two distinct molecular weight bands, differing from the corresponding band in the eluate, suggesting the presence of chemically modified, e.g., glycosylated, PDPN variants, not accessible by the used anti-PDPN antibody (Supplementary Figure S3a). Interestingly, this pattern of PDPN bands was also observed in the CD90-EVs from of the CD90-separation, hinting at a distinct PDPN^+^ EV population embedding a modified PDPN variant, without or lower CD90 co-expression (Figure 3c, d).

Validation with a pan-EV isolation kit: To validate the immunocapture approach, we applied a commercially available PAN EV kit (Miltenyi Biotec) targeting the tetraspanins CD63, CD9 and CD81. This kit is designed to isolate as much tetraspanin^+^ EVs as possible from a source population using the same principles of the CD90/PDPN separation approach. Western Blot analysis of the PAN EV separation kit using the same fibroblast EVs showed the enrichment of the targeted tetraspanin markers CD63 and CD9, along with other EV markers, in the eluate, confirming the reliability of the used immunocapture approach with our SEC-UF isolated EV source material (Supplementary Figure S3b).

In summary, we successfully separated synovial fibroblast-derived EVs into two distinct EV subpopulations using prototype anti-CD90 and anti-PDPN magnetic beads; CD90^+^ EVs reflect a CD90^high^, PDPN^high^, CD9^high^, Annexin-1^high^, CD63^low^, Flotillin-1^low^ pattern, whereas CD90-EVS display a CD90^low^, PDPN^low^, Annexin-1^low^, CD63^high^, Flotillin-1^high^ marker profile. Both populations contain equal amounts of Syntenin-1. PDPN separation yielded similar results but revealed a separate PDPN^+^ subpopulation with a potentially modified PDPN variant and lower CD90 co-expression. These findings underscore the effectiveness of our refined SEC-UF coupled immunocapture approach for isolating distinct EV subpopulations, providing critical insights into the heterogeneity of arthritis-associated EVs.

### 4.) Using the refined immunocapture approach enables the separation of CD90^+^ EVs from biofluids of higher complexity

Next, we tested the prototype bead-based immunocapture approach to target a distinct EV subset in solutions with higher EV complexity, including a mixed EV population isolated from different cell culture supernatants and, subsequentially, synovial fluids. To evaluate the robustness of our approach we conducted two different spike-in experiments under defined conditions: (i) by mixing SEC-UF isolated EVs from THP1 macrophage cell culture with SEC-UF isolated synovial fibroblast-derived EVs at a ratio of 1:1 (Figure 4a), and (ii) by spiking SEC-UF isolated synovial fibroblast-derived EVs into hyaluronidase-treated synovial fluid from arthritis patients (Figure 4d). In both experiments, we anticipated to recover a substantial portion of the spiked CD90^+^ EVs from the complex EV mixture while minimizing contamination by non-specific EV subsets or the loss of targeted CD90^+^ EVs.

**Figure 4:**
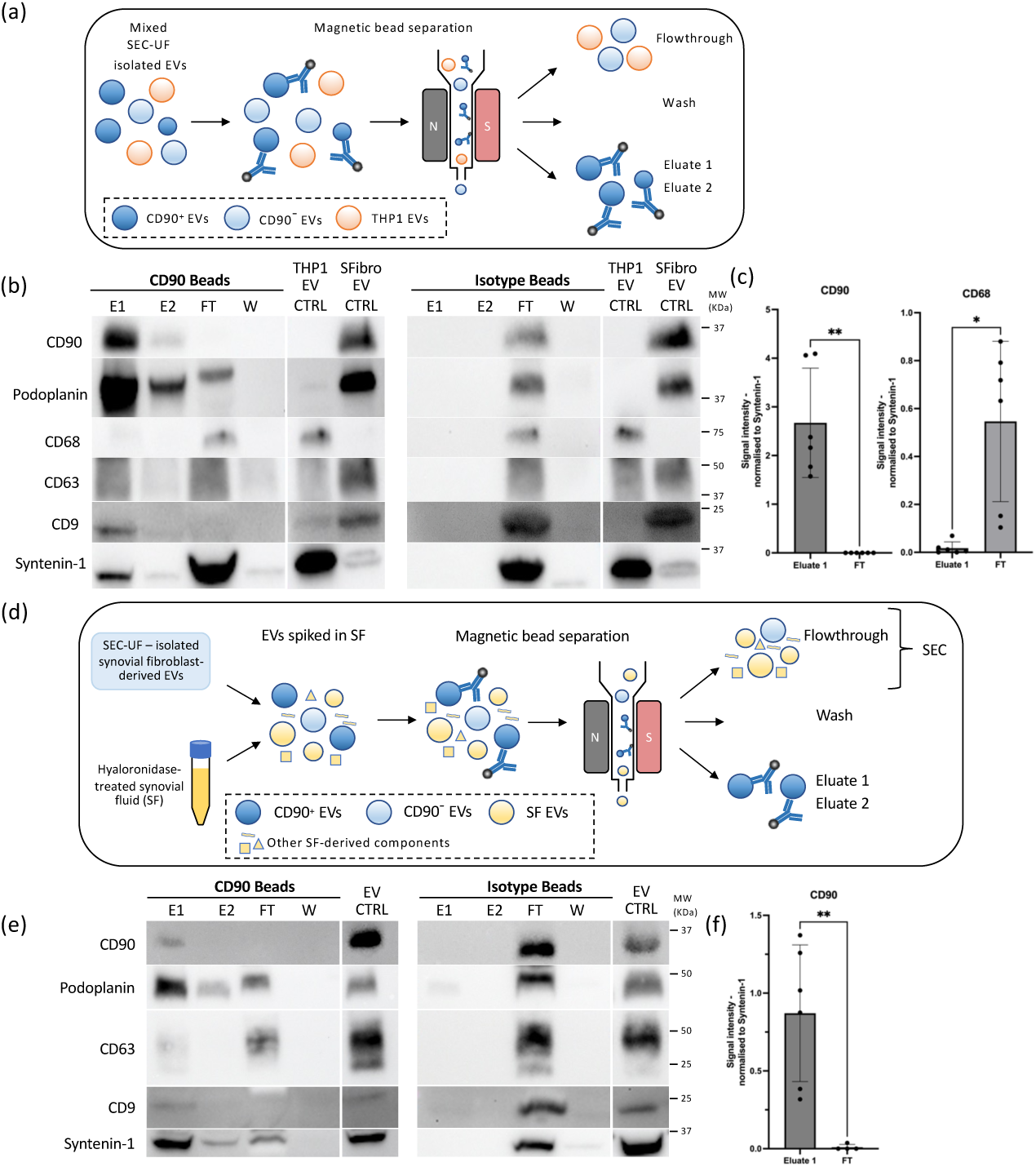
Efficient Separation of CD90^+^ EVs from complex biofluid environments using the newly established immunocapture approach. (a) Schematic representation of CD90^+^ EV separation from a mixed EV population. SEC-isolated synovial fibroblast-derived EVs (blue) were mixed at a 1:1 ratio with SEC-isolated THP1-derived EVs (orange). CD90^+^ EVs (dark blue) were selectively targeted and separated using the magnetic bead-based immunocapture approach. Antibody-bound CD90^+^ EVs were retained on the magnetic column, while unbound EVs, including CD90^−^ EVs and THP1-derived CD68^+^ EVs, were collected in the flowthrough fraction (FT). A washing step (W) was applied to remove residual unbound EVs, aggregates, and free proteins. CD90^+^ EVs were eluted from the column in two sequential elution steps (E1 and E2) to assess the elution efficiency. (b) Western Blot analysis of CD90^+^ EV enrichment and immunocapture specificity in the mixed EV population - Representative Western blot from three independent experiments showing CD90^+^ EVs enriched in the eluate (E1) and depleted in the FT. CD68, a marker for THP1-derived EVs, was detectable only in the FT and not in the E fraction. An isotype bead control confirmed the specificity of the anti-CD90 antibody, with no detectable CD90 signal in the E fraction. SEC-isolated EV input from both cell types served as a control. (c-f) Quantitative analysis of Western Blot results - Signal intensities for CD90 and CD68 were normalized to Syntenin-1, and statistical significance was assessed using student’s paired t-test. Data represent mean ± STD from three independent experiments (*p<0.05,**p < 0.005). (d) Schematic representation of CD90+ EV separation from spiked synovial fluid - SEC-isolated synovial fibroblast-derived EVs (blue) were spiked into hyaluronidase-treated synovial fluid (yellow), which encompassed no detectable native CD90 EVs. Unbound EVs, including synovial fluid-derived EVs, were collected in the flowthrough and processed via SEC to enrich EVs while excluding excess serum proteins present in synovial fluid. (e) Western Blot Analysis of CD90^+^ EV enrichment in synovial fluid spiking experiments - Representative Western blot showing CD90^+^ EV enriched in the eluate (E1) and CD90 depletion in the FT fraction. Isotype bead control confirmed the specificity of the separation, with no detectable CD90 signal in the eluate. SEC-isolated EV input served as a control for the spiked EV population.

In the first spike-in experiment, SEC-UF isolated EVs from THP1 macrophages (CD68-positive) were mixed with SEC-UF isolated synovial fibroblast-derived EVs (CD90-positive). Following bead-based separation, Western blot analysis showed a clear enrichment of CD90^+^ EVs in the eluate, with no detectable CD68 signal, indicating that macrophage-derived EVs were exclusively present in the flowthrough fraction (Figure 4b-c). This confirmed the specificity of the separation approach, effectively retrieving CD90^+^ EVs while excluding nonspecific EV subpopulations from the eluate.

The second spike-in experiment involved spiking SEC-isolated CD90^+^ synovial fibroblast-derived EVs into hyaluronidase-treated synovial fluid from arthritis patients. A synovial fluid specimen without detectable native CD90 signal was specifically chosen to ensure that any retrieved CD90^+^ EVs originated solely from the spiked population. Western blot analysis of the eluate confirmed the retrieval of CD90^+^ EVs, with no detectable signal in the flowthrough fraction (Figure 4e-f). However, the CD90 signal in the eluate was less prominent compared to the mixed cell culture experiment, suggesting reduced separation efficiency in the synovial fluid setting, potentially due to interference from synovial fluid components, such as protein aggregates or glycosaminoglycans, that may affect bead binding or separation efficacy.

In both experiments, the isotype control beads confirmed the specificity of the anti-CD90 antibody, as no CD90 signal was detected in eluates when using antibody isotype controls (Figure 4b and 4d). The separation patterns observed in these spike-in experiments mirrored those from synovial fibroblast-derived EV separation, with distinct CD90^+^ and CD90^−^ populations. Notably, the CD90− population consistently lacked CD90 enrichment, demonstrating the robustness of the approach across different experimental setups.

These spike-in experiments confirmed the capability of our bead-based immunocapture approach to separate CD90^+^ EVs from complex solutions such as synovial fluid. While separation efficiency was reduced in synovial fluid compared to a mixed cell culture setting, the method reliably enriched CD90^+^ EV subpopulations with high specificity. These findings establish the feasibility of using this refined approach for the targeted isolation of disease-relevant EV subsets from complex biofluids, paving the way for downstream applications in biomarker discovery and functional analysis.

### 5.) Isolation of synovial fluid-derived EV samples via SEC

Building on the demonstrated capacity of our bead-based immunocapture separation approach to selectively target CD90^+^ EVs in complex fluids with EV heterogeneity, we sought to isolate and characterize CD90^+^ and PDPN^+^ EVs directly from synovial fluid samples collected from arthritis patients. Due to the well-documented patient-dependent variability in EV composition within the synovial fluid, we selected two pre-screened patient samples, which displayed discrete EV patterns: one exhibiting detectable CD90 expression (Patient 5) and one without (Patient 8), as assessed by western blot in UC-derived EV isolates (Figure 1b).

Isolating EVs from synovial fluid presents significant technical challenges due to its complex molecular composition, which includes extracellular matrix (ECM) components, serum ultrafiltrate, and changing amassment of infiltrated cells, as well as its variable viscosity and other physical characteristics^62^. To overcome these challenges, we consulted the latest MISEV guidelines of the Synovial Fluid Taskforce^71^ and applied a series of pre-treatment steps adapted from protocols of published studies describing synovial fluid EV isolation via SEC^62,65^. The pre-treatment of synovial fluid is essential to avoid clotting of the SEC column and reduce contamination of co-isolated serum ultrafiltrate and ECM components. The steps included sequential centrifugation to deplete cells, precipitates and debris, as well as treatment with hyaluronidase, DNAse and protease/phosphatase inhibitors to reduce viscosity, minimize ECM stickiness and inactivate active proteases and phosphatases. Finally, the pre-treated fluid was subsequentially filtered and diluted to assure optimal performance during SEC-based EV isolation (Figure 5a, see Methods for details).

**Figure 5:**
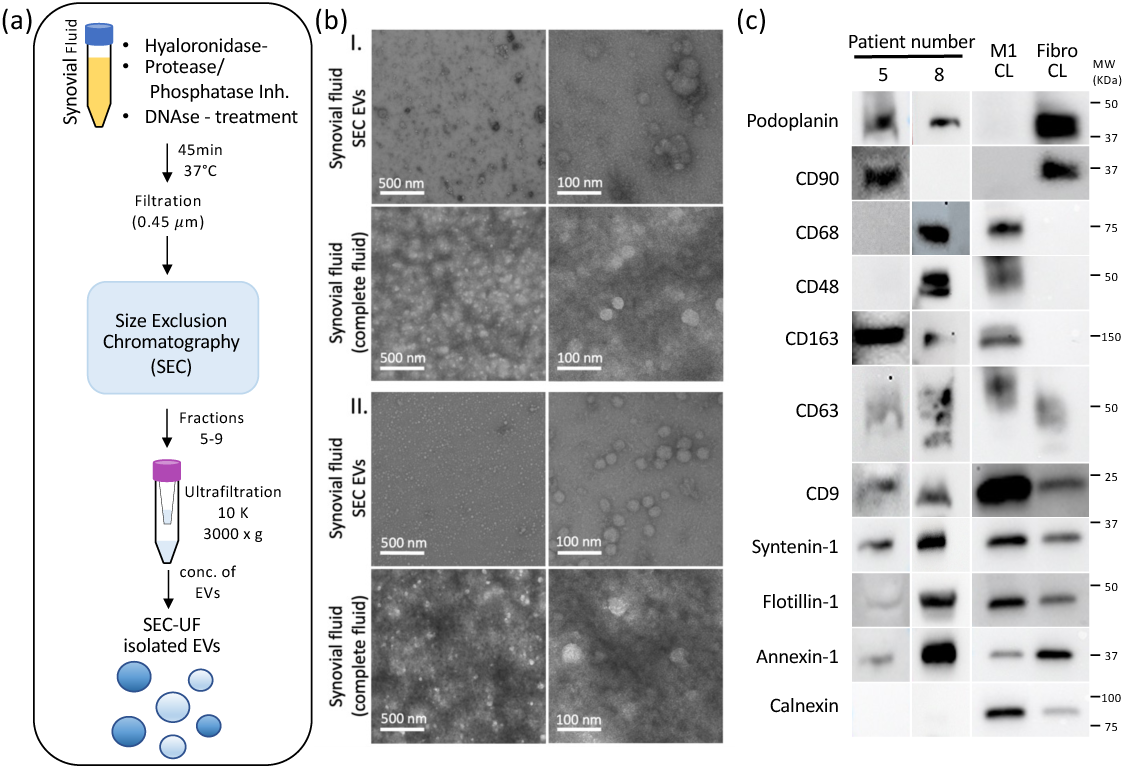
Isolation of EVs from synovial fluid via SEC shows inter-patient variability in marker expression on EVs. (a) Graphical illustration of synovial fluid-derived EV isolation via SEC. (b) TEM picture of two synovial fluid-derived EV samples isolated via SEC compared to complete synovial fluid. (c) Western Blot analysis of SEC-isolated EVs comparing two different arthritis patients.

Visualization of SEC-isolated EVs by TEM analysis revealed intact, round-shaped vesicles ranging from 30 to 200 nm, with minimal evidence of contaminating aggregates. In contrast, TEM imaging of unprocessed synovial fluid showed abundant precipitates and aggregated material, highlighting the effectiveness of the SEC-UF approach in purifying of synovial fluid-derived EVs while significantly reducing SF-derived components present the crude SF samples (Figure 5b).

Western Blot analysis of SEC-UF isolated EVs (Figure 5c) reflected the marker expression patterns observed in the ultracentrifugation pre-screening (Patient 5 and 8 – Figure 1b). CD90 expression was evident in Patient 5 but absent in Patient 8, with the reverse pattern for CD68, underscoring the inter-patient variability of synovial fluid derived EV profiles. Typical EV markers like CD63, CD9, Flotillin-1 and Annexin-1 were detected in both samples, while the EV exclusion marker Calnexin was only observed in cell lysates, confirming the purity of the isolated EV fractions.

Taken together, the pre-treatment of synovial fluid samples facilitated the successful isolation of EVs using the SEC-UF approach, significantly reducing contamination by free proteins and ECM components associated with crude synovial fluid isolates. These pre-treatment and isolation steps of synovial fluid may benefit efficient immunocapture but are crucial for downstream applications, such as LC-MS/MS-based proteomic analyses, where reducing background noise is critical.

### 6.) CD90^+^ and PDPN^+^ EV separation of synovial fluid EVs comparing SEC enrichment vs. direct biofluid approaches using the novel bead-based immunocapture method

To evaluate the necessity of SEC-UF to enrich and purify EVs from synovial fluid for subsequent immunocapture, we compared the performance of our magnetic bead-based separation method targeting CD90^+^ and PDPN^+^ EVs under two distinct conditions: (i) using pre-enriched SEC-UF-isolated synovial fluid EVs, or (ii) directly applying the immunocapture approach to pre-treated synovial fluid without SEC processing (Figure 6a). TEM imaging of both separation approaches (SEC-UF and direct) using a test sample revealed intact EVs bound to beads in the eluate fraction, while bead-unbound, free EVs accumulated in the flowthrough (Supplementary Figure S4a). Based on these results, we then performed the two distinct separation approaches for the Patient 5 and Patient 8 synovial fluid.

**Figure 6:**
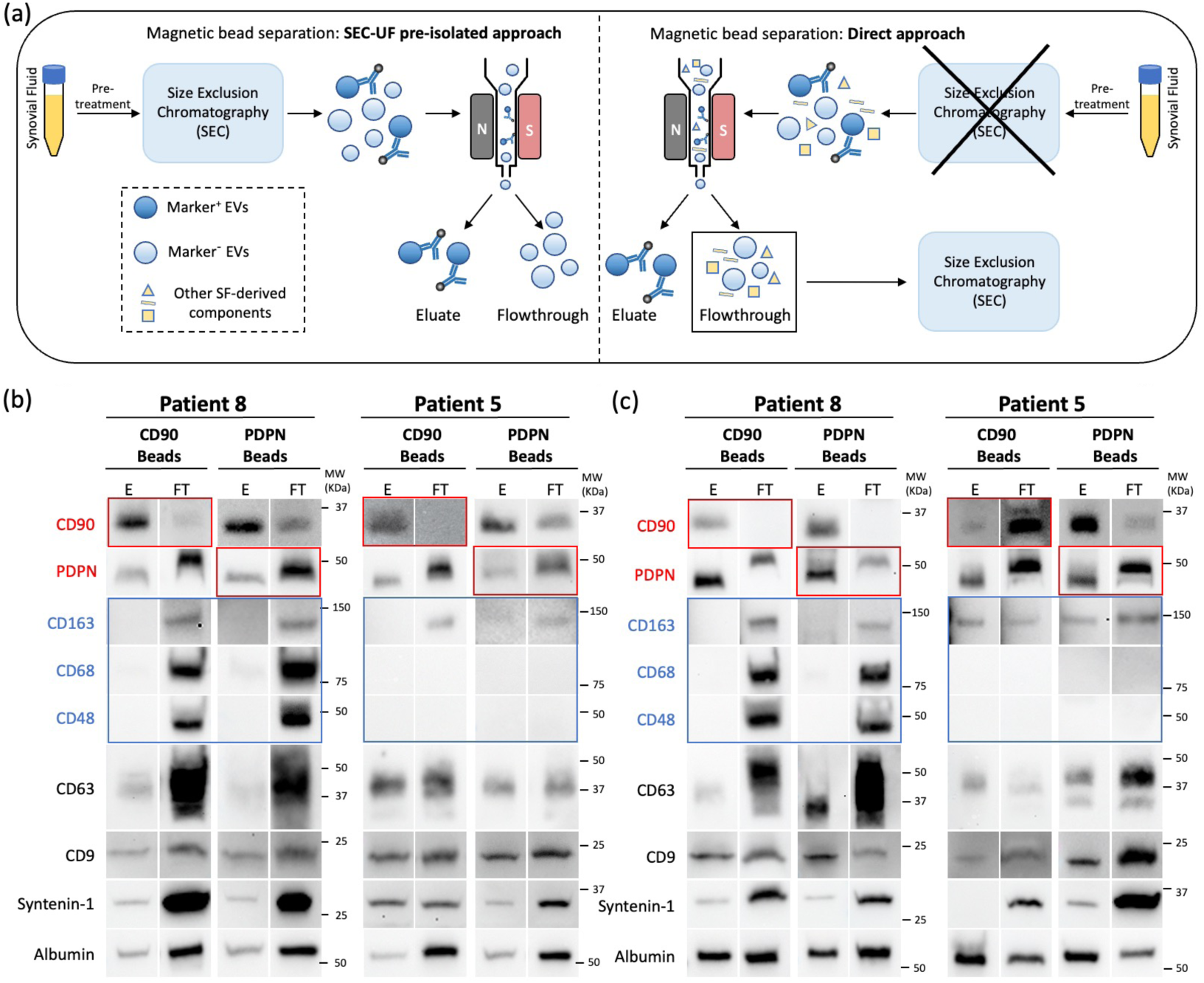
CD90 and Podoplanin MicroBead separations in synovial fluid-derived EVs. (a) Graphical illustration comparing direct usage of prototype immunocapture MicroBeads in synovial fluid with usage in pre-isolated EVs via SEC. Pre-treated synovial fluid (hyaluronidase, DNAse and phosphatase/protease inhibitor treated) was used for bead separations. FT derived from direct approach were applied to SEC following MicroBeads separation as well. (b) Western Blot analysis of prototype CD90 and PDPN MicroBead separations in synovial fluid-derived, and SEC pre-isolated EVs. Red boxes show the Western Blot signals for stromal-derived proteins (CD90 and PDPN), while blue boxes show the signals for myeloid-derived proteins (CD163, CD68, CD48). Compared are the separations from two different patients (patient 5 and 8 - see also UC-based pre-screening). (c) Western Blot analysis of prototype CD90 and PDPN MicroBead separations directly in synovial fluid of patient 5 and 8. Eluate (E), Flowthrough (FT).

We first performed the immunocapture using pre-enriched SEC-UF isolated EVs as source material for CD90 bead separation. We successfully separated a CD90^+^ EV subpopulation from Patient 5 and Patient 8 samples. Western Blot analysis reveal a clear CD90 signal in the eluate of both patients (Figure 6b), including Patient 8, where CD90 expression was undetectable in the previous analysis using a non-separated bulk EV population (Figure 1b and Figure 5c). This result demonstrated the ability of our approach to target and enrich low-abundance EV subpopulations that are otherwise undetectable in bulk EV preparations. Similar results were obtained with a separate patient sample (Patient 3, Figure 1; Supplementary Figure S4b). In correspondence with the results from the separation experiments conducted with EVs derived from cultured synovial fibroblasts, in both patients, CD90^+^/PDPN^+^ EVs were co-enriched in the eluate, while a distinct PDPN^+^ EV population with no or minimal CD90 co-expression resided in the flowthrough. This was evident from a distinct PDPN band in the flowthrough fraction running at a higher molecular weight as shown in Western blot results. Importantly, the macrophage/monocyte markers (CD163, CD68 and CD48 in Patient 8, and CD163 in Patient 5) were exclusively present in the flowthrough, confirming the specificity of the CD90 bead separation to target stromal cell type-derived EVs. The tetraspanin markers (CD63 and CD9) displayed similar distribution profiles as observed in earlier experiments, while elevated Syntenin-1 and albumin signals in the flowthrough reflected the untargeted EV population and residual free serum proteins still present following the SEC-UF isolation. A parallel PDPN-bead-based separation yielded comparable results, though enrichment for the CD90^+^/PDPN^+^ co-expressing EVs was less pronounced in the eluate (Figure 6b).

We next applied the direct immunocapture from pre-treated synovial fluid (Figure 6c). In the direct approach without SEC-UF enrichment, the bead-based separation was first validated using concentrated supernatant of cultured synovial fibroblasts, which displayed results consistent with those obtained from the SEC-UF-isolated EVs of cultured synovial fibroblasts (Supplementary Figure S4c, compare Figure 3). Then, we performed the immunocapture approach directly in synovial fluid from Patient 5 and Patient 8, pretreated as described, but without the SEC-UF step. As the flowthrough in direct approach still included the neat synovial fluid components at a high proportion of total protein, that fraction needed to be processed via separate SEC-UF to enable the detection of EV-derived proteins (Figure 6a, right panel). Western blot analysis of the direct approach displayed different results in Patient 8 compared to Patient 5 synovial fluid (Figure 6c). The CD90- and PDPN-bead separation worked successfully in Patient 8, showing a similar profile for the tested markers as observed in the approach with SEC-UF isolated EVs. Western blot analysis showed successful separation of an enriched CD90^+^/PDPN^+^ EV subpopulation in the eluate, while CD90^-^/PDPN^+^ EVs and the macrophage marker-positive EV populations were confined to the flowthrough fraction (Figure 6c, left panel). However, higher albumin levels in the eluates of the direct vs SEC-UF approach indicated remaining serum protein from the crude synovial fluid, likely due to the absence of the SEC-UF purification step. This observation aligns with the known formation of a protein corona on the surface of biofluid-derived EVs^72^.

In contrast, the direct CD90-bead separation approach failed to enrich CD90^+^ EVs in the Patient 5 sample (Figure 6c, right panel). No CD90 enrichment was observed in the eluate, and the macrophage marker CD163 was not effectively separated between fractions. The direct PDPN-bead separation, however, was successful for Patient 5, producing results consistent with those obtained in the SEC-UF-enriched approach as before. These findings highlight the variable performance of EV immunocapture methods when applied directly to unprocessed biofluids, in line with previous reports suggesting that factors within crude biofluids such as plasma can interfere with immunocapture efficacy^73^.

Our data demonstrated that the novel bead-based immunocapture approach reliably enriches CD90^+^ and PDPN^+^ EV populations from SEC-UF isolated synovial fluid EVs with high specificity. The enrichment of stromal-derived EVs (CD90^+^/PDPN^+^) was validated by the absence of macrophage markers (CD163, CD68, CD48) in the eluate, which were confined to the flowthrough. In the direct approach, while CD90^+^/PDPN^+^ EV separation was feasible in some samples (Patient 8), variability in outcomes, likely due to patient-specific factors present in the crude synovial fluid, reduced the reliability of the method. This was particularly evident in the failure to enrich CD90^+^ EVs from Patient 5 synovial fluid, where direct binding efficiency of the bead-coupled anti-CD90 antibody appeared to be compromised. These findings underscore the importance of pre-enriching and purifying EVs from synovial fluid, e.g., via a SEC-UF, to minimize contamination with secondarily attached serum proteins and improve immunocapture reliability. This is especially important for sensitive downstream analysis like LC MS/MS that are suffering from high range of abundant proteins. The direct approach, while promising, remains subject to unpredictable outcomes due to the complex nature of crude biofluids and patient variability and warrants further optimization to enhance its consistency and reliability for downstream analyses.

### 7.) Differential expression analysis of MS-measured proteins in separated CD90^+^ and CD90^-^ EVs from synovial fluid

Mass spectrometry-based proteomics, such as LC-MS/MS, is increasingly recognized as the gold standard for EV protein profiling offering unparalleled sensitivity and specificity in identifying complex protein signatures^74^. We therefore applied comprehensive MS-based proteomic profiling with the means to validate the novel bead-based immunocapture separation approach for isolating cell type-specific small EVs from synovial fluid of arthritis patients. In accordance with previous studies profiling arthritic synovial fluid EVs, we used highly purified SEC-UF EVs as input material for LC-MS/MS analysis^65^. This approach was previously demonstrated to overcome the technical limitations of MS-based proteomics imposed by unfractionated biological samples with a wide dynamic range in their protein concentrations^75^. This is especially an issue in a complex samples such as synovial fluid, in which high-abundance plasma proteins from serum infiltrates and ECM components can mask and suppress the identification of low abundance EV-specific proteins, potentially obscuring critical information on subtle, disease-relevant components of the EV proteome^62^. By coupling SEC-UF EV enrichment with an immunocapture step, our workflow aimed to increase the EV purity, thereby enhancing the detection of EV-specific proteins beyond what has been published in previous studies assessing synovial fluid EVs. Indeed, the MS analysis of SEC-UF isolated and CD90 ^+^/^-^ separated SF EVs from arthritis patients (n=11) resulted in a high-resolution dataset of 3306 identified proteins). To assess the coverage of our SF-EV proteome in relation to (i) small EVs in general, and (ii) to estimate the proportion of yet unidentified proteins, which are relevant for synovial fluid but excluding the portion of plasma-attributed proteins, we compared our data against two reference data sets (Supplementary Figure S5a). These included a curated set of 7862 human MS-measured small EV proteins cataloged in ExoCarta database, a publicly accessible curated repository compiling protein/RNA/lipid cargo data identified in small EVs from > 1200 conducted studies^76^ (Version 6, 2024), and a set of 551 high-abundance plasma proteins (concentration > 0.1mg/L assessed in human blood plasma) listed in the Human Protein Atlas database (HPA/PeptideAtlas). This analysis revealed that roughly 90% of the EV proteins in our dataset are listed in the set of small EV proteins in ExoCarta (2961/3306), confirming a clear enrichment of EV proteins. Further, 74% of the 551 most abundant plasma proteins (405/551) were present in our SF-EV proteome, highlighting the substantial contribution of serum infiltrates in synovial fluid. Still, our refined SEC-UF EV isolation strategy enabled the detection of 246 additional EV proteins not previously reported in ExoCarta, likely representing low-abundance proteins related to EV populations present in arthritic synovial fluid that were previously undetectable in bulk SF-EV proteomic measurements.

Next, we evaluated our established immunocapture separation approach targeting stromal-derived EV populations from synovial fluid of arthritis patients (n= 11) by performing a bioinformatic analysis of the MS-measured proteins present in the two separated fractions, “Eluate (E, CD90^+^ EVs)” and “Flowthrough (FT, CD90^-^ EVs)”. A principal component analysis (PCA) of the MS data clearly visualized the separated clusters of the E *vs* FT fraction samples. Still, the complex proteomic landscape of synovial fluid EVs was evident in the PCA, as the difference between E *vs* FT was less pronounced compared to the corresponding fraction samples assessed in a parallel MS-measured EV set derived from *ex vivo* cultured synovial fibroblast cells (Supplementary Figure S5c). This apparent narrower clustering of E and FT fraction samples in the SF-EVs was likely driven by the content of absorbed synovial fluid proteins equally present on EVs in both fractions.

Next, we profiled the identified SF-EV protein sets in the two separated fractions in detail by differential expression analysis. This analysis found a total of 616 proteins significantly differentially expressed (adjusted p-value < 0.05). To allocate the significant proteins to the two separated fraction E and FT, we set a cut-off at log2FC = +/- 1.5, resulting in 269 proteins significantly enriched within the CD90**^+^** EVs in fraction E, compared to 296 proteins enriched in FT, while 51 proteins were below the set cut-off and designated as significant but shared (S) (Figure 7a, Supplementary Figure 5b). The first inspection of the enriched proteins confirmed the enrichment of CD90/THY1 in fraction E, along with further stromal-associated proteins, like ADAMTS7, FAP, PDGFRB and EGFR. In contrast, proteins like CXCR4, CD8A, CD48 and MARCO, all markers for myeloid and lymphoid cell types, were enriched in the FT fraction (Figure 7b).

**Figure 7:**
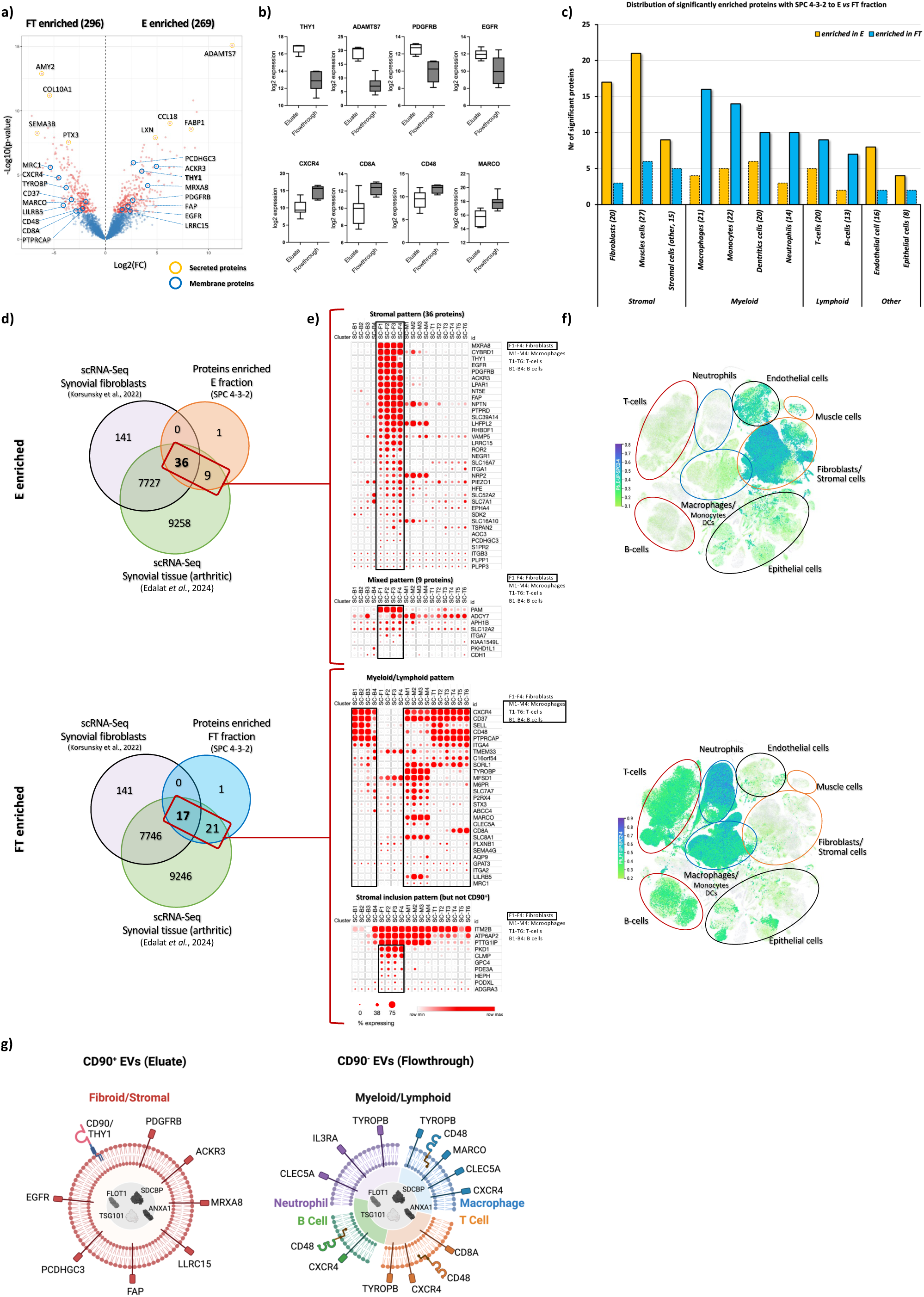
Differential expression analysis of SF EV proteome eluate (E) vs flowthrough (FT) fraction confirms successful separation of fibroblast/stromal-derived EVs from synovial fluid of arthritis patients using the prototype CD90-Microbead immunocapture approach. (a) Volcano Plot displaying differentially expressed SF-EV proteins (3306) in the CD90^+^ EV fraction (Eluate) versus the non-captured EV fraction (Flowthrough). Proteins significantly enriched in the Eluate fraction samples are plotted as red dots, while proteins with no significant differential expression are depicted as blue dots. Proteins of interest are highlighted with colored circles, indicating membrane (blue circles) or secreted proteins (orange circles). (b) Violin blots displaying Log2 expression values for four proteins significantly enriched in E fraction samples (THY1, ADAMTS17, PDGFRB, EGFR) and four proteins significantly enriched in FT fraction samples (CXCR4, CD8A, CD48, MARCO). (c) Cell Type Signature (CTS) distribution of differentially enriched proteins. Bar graph shows the number of significantly enriched surface proteins (SPC score 4-3-2) across the 11 curated CTS groups in E *vs* FT fraction, highlighting their cell type-specific association. (d) Comparison of differentially enriched SF-EV proteins with scRNA-Seq data from synovial tissue. Venn diagram illustrating the overlap between SF EV proteins with SPC score 4-3-2 significantly enriched in E or FT fraction and genes signatures identified in synovial fibroblasts (Korsunsky *et al.*, 2022^79^) or synovial tissue cell populations in inflammatory arthritis (Edalat *et al.*, 2024^80^). The intersections classify E-enriched proteins as predominantly stromal-derived, while FT-enriched proteins overlap with multiple cell populations present in arthritic synovium, including immune cells. Projection of E- and FT-enriched proteins onto two independent single cell atlases from available as online portals, (e) the Broad Institute Single Cell Portal (SCP, https://singlecell.broadinstitute.org/single_cell) and (f) Tabula Sapiens (https://tabula-sapiens.sf.czbiohub.org). SF-EV proteins in E or FT located in the intersections in figure d, were projected onto the two single cell transcriptomic atlases. SCP projection of the selected enriched protein set using the Accelerating Medicines Partnership (AMP) Phase I project RA dataset (Zhang *et al.*, 2019^39^) reveals a dominant stromal expression pattern for E-enriched proteins compared to a primarily myeloid/lymphoid expression pattern with minor stromal inclusions in FT-enriched proteins, as visualized by expression dot blots. Visualization by UMAP clustering of E *vs* FT enriched proteins using the Tabula Sapiens Single Cell Atlas showed a similar stromal-dominated expression pattern in the E fraction and a myeloid/lymphoid-skewed signature in the FT fraction. Circle colors indicate different cell types, lymphoid (red), myeloid (blue), stromal/fibroid (orange) and other cell types (black). (g) Overview of CD90^+^ separated EVs in Eluate with stromal membrane markers *vs* EVs in the Flowthrough with myeloid/lymphoid markers. Schematic representation summarizing the findings, showing that CD90+ EVs enriched in the eluate contain stromal membrane markers, whereas non-captured FT-derived EVs predominantly harbor myeloid/lymphoid markers, confirming the successful immunocapture-based separation of distinct synovial EV subpopulations. This part (g) was created with Biorender.com.

The amassed presence of activated immune cells in the synovium during synovitis was reflected by the remarkable proportion of CD markers identified in our SF-EV proteins. A set of 150 out of 384 listed CD markers (7= significant in E fraction, 10= significant in FT fraction) were detected thereby convincingly reflecting the heterogenous nature of EV populations present in the synovial fluid of arthritis patients. In analogy to CTAPs assessed in recent scRNA-Seq studies^40^, we allocated the identified CD markers to a myeloid, lymphoid and stromal cell continuum (Table 1 CD markers). To further explore the heterogeneity of the synovial EV population and identify additional cell type-specific markers in our E or FT fraction in addition to CD markers, we curated a panel of 10 defined cell type signatures (CTS) representing cell types relevant in synovitis (gene sets based on the HPA database and PangLaoDB for selected cell types). We selected 9 cell types representing the myeloid-lymphoid-stromal cell continuum according to published scRNA-Seq data profiling synovial tissue cell populations. The curated 9 CTS included protein-coding genes attributed to macrophages/monocytes, dendritic cells and granulocytes (neutrophils) for the myeloid group; T-cells (including NK-cells) and B-cells for the lymphoid group, while fibroblasts, muscles cells and a combined set of stromal-related cell types (adipocytes, chondrocytes, osteoblasts and osteocytes) were comprising the stromal group. Finally, we also added an additional CTS representing stromal-associated endothelial cells, as the vasculature surrounded by mural cells/pericytes plays a significant role in arthritis pathology ^45^.

**Table 1:** Examples of 150 CD markers identified in synovial fluid-derived EVs of arthritis patients classified according to their cell type association.

To exploit cell type-specific markers on EVs as potential novel targets for separation approaches, they must be exposed on the EV membrane accessible for immunocapture. For identifying such markers located on the EV surface, we applied the cell surface protein consensus (SPC) score ranging from 0-4, which predicts the likeliness of a protein to be localized to the cell membrane by using a curated surfaceome dataset^77,78^. Scores 4-3-2 classify “cell surface location”, while score 1 refers to “predicted localization” and 0 to “not localized” to cell surface. Scoring our EV proteome according to SPC, we found 559 proteins including 90 (46 in fraction E, 39 in fraction FT, 5 in shared (S)) significantly enriched proteins with SPC score 4-3-2.

We then cross-referenced significantly enriched proteins with the SPC scores 4-3-2 with the curated 10 CTS, which confirmed a stromal/fibroid signature in the E fraction, with matching proteins from CTS groups for fibroblasts, muscle cells and further stromal cells, all enriching with CD90**^+^** separated EVs, while the myeloid and lymphoid CTS were enriched in the FT fraction (Figure 7c). Interestingly, the separate CTS designated to an endothelial cell type also showed an enrichment in the E fraction, indicating EVs from endothelium-associated cells, such as pericytes present in synovial tissue, which express CD90 and share a mixed stromal-endothelial signature^45^. Cells resident in a given tissue-environment exhibit tissue-specific expression patterns, which can either partially overlap with the expression pattern of a different cell type present in the same niche, or can differ significantly between distinct tissues^79^. To validate the detected stromal markers, which were co-enriched on the separated CD90**^+^** EVs in fraction E, with synovial tissue specific cell population signatures, we compared the E *vs* FT proteins with recently published scRNA-Seq data profiling synovial fibroblasts^79^ or a single cell atlas of synovial tissue in inflammatory arthritis^80^ (Figure 7d, Supplementary Figure S6a). The comparison of the 46 proteins with SPC score 4-3-2 that were significantly enriched in the E fraction, revealed a prominent overlap with the synovial fibroblast data set (78 %, 36 proteins), including CD90/THY1, ACKR3, EGFR, FAP, LRRC15, MXRA8, NRP2, PCDHGC3 and PDGFRB. Interestingly, the enriched set also included proteins not formerly associated to a stromal cell type such as EPHA4, LPAR1, MELTF, NPTN, NT5E, PTPRD and ROR2. In contrast to the prominent stromal signature in E, the corresponding 39 proteins enriched in FT showed only a 44 % overlap (17 proteins), while 21 proteins not overlapping with the synovial fibroblast data set included markers generally assigned to either T-cells, B-cells macrophages, monocytes or neutrophils, including CD8A, PTPRCAP, CXCR4, MRC1, CD48, CD37, LILRB5, M6PR, MFSD1, MRP4/ABCC4, SLC7A7 or CLEC5A. This difference in E *vs* FT attributed cellular profile underlined the role of CD90/THY1 to discriminate myeloid and lymphoid cell types. However, the FT fraction still included proteins associated or shared with stromal cells, which were represented by the 17 proteins in the synovial fibroblast intersection, potentially representing CD90-negative stromal/fibroblast-derived EVs. These included CLMP, PODXL, PDE3A, HEPH, ADGRA3 or GPC4 with no clear myeloid or lymphoid association. The FT fraction also included the immune cell-related proteins MARCO, TYROBP, SELL and AQP9, which were also expressed by synovial fibroblasts. Several of the remaining significant proteins in E and FT without clear cell type signature represented vesicular proteins such as TSPAN2, VAMP5, RHBDF1, STX3 or SORL1. Notably, most of the intrinsic EV proteins, including ANXA1, FLOT-1, TSG101 and SDCBP or the mentioned plasma-derived proteins were not enriched in either fraction, but were shared and located in the pool of not-significantly enriched proteins.

Finally, we used two different web-based single cell gene expression portals to visualize the expression pattern of our enriched proteins in fraction E *vs* FT on a cell type-specific or a global tissue-specific level. We used the Broad Institute Single Cell Portal (SCP) with the implemented data set from the Accelerating Medicines Partnership (AMP) Phase I project (data set 1 Rheumatoid Arthritis^39,81^ to represent cells of the synovial lining and sublining in inflammatory arthritis. To assess the expression profile of our enriched proteins on a global level we used the human cell atlas provided by Tabula Sapiens, which provided expression data of 1.1x 10^6^ cells derived from 28 organs of 24 normal human subjects^82^. Projection of the significant proteins enriched in fraction E on the synovial cell atlas revealed a strong fibroblast/stromal expression pattern, while the significantly enriched proteins in fraction FT exhibited a mixed myeloid/lymphoid pattern with some stromal inclusions (Figure 7e, Supplementary Figure S6 a-c). A very similar expression pattern was shown for the projected E *vs* FT proteins on the Tabula Sapiens global cell atlas (Figure 7f). This pattern was also confirmed by extracting the quantified expression values of selected cell types from the CZ CELLxGENE portal, which were comparable to the 10 CTS generated in the current study, and plotting them on a radar distribution blot (Supplementary Figure S6c). In conclusion, the proteomic and different bioinformatic validation assessments confirmed the successful EV separation method based on our established CD90^+^ immunocapture approach to target a stromal EV subset from the heterogenous EV population present in synovial fluid of arthritis patients (Figure 7g).

## Discussion

Extracellular vesicles (EV) have emerged as promising biomarkers for the diagnosis, prognosis, and monitoring of therapeutic responses across various diseases. Emerging data suggests that EVs in arthritis harbor disease-specific characteristics which are contributing to disease progression by modulating pathogenetically relevant molecular pathways through their specific cargo^11,59,65,83^. This highlights the potential of EVs not just as biomarkers, but also as key players in disease pathogenesis, opening avenues for novel therapeutic interventions^84^. Consequently, the presence of EV populations in biofluids, along with their specific cargo, offer a window into the pathophysiological state of the cells and tissue from where they emanate. This is particularly relevant in the context of arthritis, especially in RA, where the cellular basis of the underlying chronic synovitis is intricate and manifests as a variety of diverse histopathological phenotypes tied to distinct cellular populations in the synovial membrane^40^. Recent strides in single-cell omics technology and bioinformatics have unveiled novel functional subsets of tissue-resident synovial fibroblasts, macrophages, and infiltrated immune cells, each playing unique roles in RA pathogenesis and potentially also the treatment response^66,85^. The ability to identify distinct EV populations in liquid biopsies from synovial fluid or plasma that reflect the presence and activity of disease-relevant synovial tissue cells would provide a less invasive alternative approach to tissue biopsies. We therefore aimed at developing a method to separate cell-type specific EVs from synovial fluid allowing the subsequent analysis of their proteomic signatures. Herein we show a refined EV separation approach to target and analyse arthritis-specific EV subpopulations from cultured *ex vivo* synovial fibroblasts or from patient-derived synovial fluid using prototype nano-sized magnetic beads. To the best of our knowledge, we present the first study demonstrating magnetic bead-based immunocapture of cell-type specific EVs, either coupled to a SEC-purification pre-enrichment step, or directly from preprocessed synovial fluid.

However, the suggested use of EVs in biomarker discovery for arthritis is still hampered due to considerable technical challenges involving the isolation and analysis of distinct EV subsets from patient-derived synovial fluid. First, the EV pool present in synovial fluid is a highly heterogenous mixture of different EV populations, which originates from cells in the surrounding periarticular tissue as well as infiltrating immune cells^11,58^, making it difficult to define separate disease-relevant EV subsets by using classical bulk EV assessments. The issue of heterogeneity is further complicated by a considerable patient variability, reflected by changing composition of the inherent EV pool or the chemo-physical parameters of the synovial fluid^60^, altogether limiting precise and standardized isolation methodologies. Moreover, synovial fluid is a protein-rich lubricant that exhibits a unique ECM composition and, in the case of active arthritis, is enriched with secretions of immune cells and high-abundance plasma proteins^86^. Especially plasma proteins and ECM fragments constitute a substantial portion of total protein in isolated synovial fluid EVs^72,87^, thereby representing a critical factor for downstream EV proteomic profiling by increasing the risk to obscure low-abundance proteins of interest. This is especially an issue in MS-based proteomics, which exhibits sensitivity limitations when measuring samples with a wide dynamic range in protein concentration^88^. Hyaluronidase treatment of cell free synovial fluid before EV isolation was shown to reduce viscosity and facilitate EV isolation^87^. Other strategies to restrict the amount of non-nascent EV proteins secondarily adsorbed to the surface include proteinase K treatment to strip off “contaminating” proteins from the surface of isolated EVs^62^, or processing the EVs by selective affinity-base purification steps targeting IgG or proteoglycans^89^. Though, these purification strategies reduce the portion of adsorbed proteins, they can lead to decreased EV yield, remove essential surface proteins and destroy the protein layer that constitute the so-called corona present on biofluid-derived EVs^72,90^. Especially in immune cell-derived EVs the corona might play a critical functional role and loss of the corona is leading to reduced uptake and disrupted interaction with immune cells ^91–93^. Thus, the surface of (small) EVs was considered as primary platform for functional interactions in immune modulation^94^, acting in paracrine/autocrine signalling based on their exposed receptors and ligands. The surface is also serving as antigen-presenting hub by exhibiting MHC I/II and Fcgamma receptors, coupled immunoglobulins, as well as cell adhesion components, which absorb DAMPs and active matrix modulation proteins (OA) or posttranslationally modified proteins relevant in autoimmune diseases such as RA (e.g., citrullinated vimentin)^19,95^. Therefore, a targeted EV isolation strategy directly from synovial fluid, as demonstrated by our direct immunocapture approach using prototype nanosized magnetic CD90/PDPN-beads, might be essential to preserve the protein corona and to investigate the functional role of synovial fluid EVs in downstream immune-modulation experiments.

To achieve a high-resolution profile of EV cargo and EV surface markers, we applied our established fit-for-purpose approach coupling SEC-UF EV isolation with subsequent immunocapture of a selected EV subset. This refined isolation process including gravitational washing during SEC and magnetic bead separation is aimed to gently purify the EV surface from loosely-attached free proteins or protein-aggregates without disrupting the corona. Indeed, this approach provided in a comprehensive proteome, enabling the identification of membrane-associated proteins, like immune-cell-related receptors, integrins and cell adhesion proteins. Still, proteins classified as secreted were also detectable, which are either bound to the EV surface or packed as intraluminal cargo. In contrast to conventional SEC-EV isolations, a SEC-coupled immunocapture approach unites both, high purity EV isolation with the possibility for in depth downstream analysis of a targeted EV subpopulation. Immunocapture is a classical approach used in molecular biology for cell-separation relying on magnetic-beads coupled to antibodies targeting a specific antigen present on a cell or more recently a vesicle. The available beads used in previous EV-studies highly differ in their size and composition, which influence immune capture efficiency, unspecific binding of proteins and downstream experiments. Typically used for EV isolations are magnetic Dynabeads (size: 2800 - 4500 nm)^96^, Exo-CAP Streptavidin Beads (3000 nm)^97^, Pierce Protein A Magnetic Beads (1000 nm)^98^ and Microbeads (50 nm)^99–101^. For our immunocapture approach, we also applied the Microbeads technology, which offers the advantage that the nano-sized beads remain soluble and homogeneously dispersed in the reaction solution thereby allowing an efficient EV-to-antibody interaction.

Bead separations have been reported previously as an alternative EV isolation method for UC, precipitation or SEC using beads targeting tetraspanins (CD63, CD9, CD81)^98,102–104^ or other abundant molecules exposed on the EV surface like phophatidylserine^105^, for isolating an entire EV population (bulk EVs) from cell cultured supernatants or biofluids like plasma. Other studies used beads to separate a specific disease-related EV subpopulation e.g., Pfeiffer et al. have used anti-KIT antibody-coupled Dynabeads from cell cultured human mast cell-derived EVs to investigate a KIT^+^ EV subpopulation^96^. Additionally, Dynabeads were used to separate NRXN3^+^ EVs from CSF^106^ or CD63^+^, CD9^+^ and L1CAM^+^ EVs from plasma^107^. Another study used L1CAM-based immunocapture to separate neural-enriched EVs investigating the miRNA cargo for differences between ALS patients and controls^108^. Further, several studies used EpCAM1 MicroBeads to investigate epithelial cell-derived EVs in regard of cancer investigations^100,109,110^.

Our EV separation approach was targeted against CD90/THY1^+^ EVs derived from CD90/THY1^+^ fibroblasts. These fibroblasts were reported previously for their expansion and involvement in inflammation in the synovium from patients with RA^39,47^. The transfer of FAPα^+^ THY1^+^ fibroblasts in mice resulted in a more severe and persistent inflammatory arthritis compared to FAPα^+^ THY1^-^ fibroblasts^44^. Moreover, single cell transcriptomics of RA synovial tissue identified seven different subsets of fibroblasts describing the THY1^+^ CDH11^+^ PDPN^+^ CD34^-^ subset as inflammatory subset highly proliferating, expressing pro-inflammatory cytokines and showing invading cell characteristics in vitro^47^. EVs derived from this cell population might mirror the inflammatory nature of their originating cells, by e.g., transferring inflammatory cargoes and might represent an interesting disease-specific subpopulation of EVs.

By analyzing the protein composition of the separated EV subsets using MS-based proteomic profiling, we aimed to confirm the successful separation of a CD90/THY1^+^ EV subset in fraction E, but also reveal cell-type-specific protein signatures enriched in the FT fraction. To identify the cellular origin of these separated EV subsets in E vs FT, we performed a cell origin profiling strategy by using a combinatorial bioinformatic approach. We first focused especially on membrane-located proteins identified in our dataset by using the cell surface consensus (SPC) score informing about the likeliness of a protein to be located on the exofacial side of the cell membrane. In a second step, we identified stromal, myeloid and lymphoid cell-type signatures by cross-referencing the selected membrane-associated proteins with curated cell-type specific profiles relevant for synovitis using public databases (e.g., HPA and PanglaoDB), and single-cell RNA sequencing (scRNA-Seq) atlases such as Tabula Sapiens. Finally, to confidently infer the stromal identity of the CD90/THY1^+^ enriched protein with regard to synovial tissue, we projected our EV proteomic profiles of E vs FT on a scRNA-Seq data set of arthritis-derived synovial tissue. This analysis confirmed the stromal synovial tissue-derived profile of the separated CD90/THY1^+^ EVs and revealed several co-enriched proteins with reported contributions in synovitis in RA and OA. For instance, the proteinase ADAMTS7 and the serine protease FAP both are overexpressed in RA and OA promoting the breakdown of ECM^111–113^. Additionally, PDGFRB - a part of the platelet-derived growth factor (PDGF) receptor - was detected on CD90/THY1^+^ EVs. This receptor is expressed at increased levels in RA and OA synovial tissues and its activation is involved in invadosome and aberrant bone formations^114–116^.

Finally, we identified EGFR co-enriched on CD90/THY1^+^ EVs, known for its increased expression in RA patients compared to healthy controls. EGFR was reported as a therapeutic target in RA as it has been shown for its involvement in synovial fibroblast proliferation and induction of cytokine expression. Application of cetuximab, a mAb against EGFR, could ameliorated RA symptoms in patients^117,118^. Noticeably, in contrast to that, the overexpression of EGFR in mice showed therapeutic effects for OA^119^, highlighting the different disease patho-mechanisms underlying RA and OA.

In summary, we validated the successful separation of a CD90/THY1^+^ stromal-derived EV subpopulation by using MS-based proteomic analysis and by projecting the identified proteins to scRNA-Seq atlases of synovial cells and tissue. This refined SEC-UF-coupled immunocapture approach allows to investigate the distinct proteomic signatures of separated disease-relevant EV subsets derived from the synovium of patients with different forms of arthritis. The proteomic profiling of separated EV subsets may provide insights into disease pathogenesis and lead to the discovery of novel biomarkers supporting diagnosis, monitoring, and guiding personalized treatment strategies in arthropathies.

## Methods

### Cell culture

Synovial fibroblasts were isolated from RA- and OA-patient tissues received from the Department of Orthopedics and Department of Rheumatology University hospital Basel (ethics review board approval EKNZ 2019-01693). In addition, the THP1 cell line (WT) from Sigma-Aldrich was used.

Cells were maintained in an incubator at 37 °C (5 % CO_2_) and cultivated in T175 (synovial fibroblasts) or T75 (THP1) flasks in RPMI-1640 medium supplemented with 10 % FCS (Gibco), 1 % Penicillin/Streptomycin and 25 mM HEPES. Synovial fibroblasts were passaged all 10-14 days depending on cell density, while THP1 were passaged, when cells reached a density of 1x10^6^ cells/ml. For all experiments, both cell types were used during passages 2-10

### Synovial fluid and synovial fibroblast samples: Patient information

| | Disease | Treatment | RF | Anti-CCP | CRP | White cell count (in cells/μl) | DAS28 | Age ( years) | Gender | Note | Used in |
| --- | --- | --- | --- | --- | --- | --- | --- | --- | --- | --- | --- |
| 1 | Rheumatoid Arthritis | No treatment | negative | positive | 5.3 | 19400 | 2.17 | 81 | Female |  | UC, MS |
| 2 | Rheumatoid Arthritis | Cimzia | positive | positive | 23.5 | 8120 | 4.11 | 48 | Female | also Psoriasis | UC |
| 3 | Rheumatoid Arthritis | No treatment | positive | negative | NS | 5444 | NS | 84 | Female |  | UC |
| 4 | Rheumatoid Arthritis | No treatment | negative | negative | 27 | 28530 | 3.52 | 81 | Female | Acute arthritis | UC, MS |
| 5 | Rheumatoid Arthritis | JAK | positive | positive | 1.3 | 210 | 4.17 | 81 | Female | Not inflamed puctate | UC, MS,WB |
| 6 | Rheumatoid Arthritis | JAK | positive | positive | <0.3 | 240 | 2.07 | 69 | Female | Remission RA | UC |
| 7 | CPPD | Metothrexat | negative | negative | NS | 3590 | NS | 58 | Male |  | UC |
| 8 | CPPD | No treatment | negative | negative | 38.1 | 21900 | NS | 76 | Female |  | UC,WB |
| 9 | CPPD | NS | NS | NS | NS | NS | NS | NS | NS |  | UC |
| 10 | CPPD | NS | NS | NS | NS | NS | NS | NS | NS |  | UC |
| 11 | Gout | No treatment | negative | neagitive | 124.1 | 15900 | NS | 89 | Male |  | UC |
| 12 | Osteoarthritis | NS | NS | NS | NS | NS | NS | 90 | Female |  | UC |
| 13 | Osteoarthritis | NS | NS | NS | NS | NS | NS | 77 | Female |  | UC |
| 14 | Osteoarthritis | NS | NS | NS | NS | NS | NS | 69 | Female |  | MS |
| 15 | Osteoarthritis | NS | NS | NS | NS | NS | NS | 71 | Male |  | MS |
| 16 | Osteoarthritis | NS | NS | NS | NS | NS | NS | 67 | Male |  | MS |
| 17 | Osteoarthritis | NS | NS | NS | NS | NS | NS | 72 | Female |  | MS |
| 18 | Osteoarthritis | NS | NS | NS | NS | NS | NS | 70 | Female | Presence of Synovial inflammation, hypertrophy, bone cysts and osteophytes | MS |
| 19 | Inflammatory Arthritis | No treatment | NS | NS | 2.6 | NS | NS | 66 | Male | Knee | MS |
| 20 | Rheumatoid Arthritis | No treatment | positive | positive | 6.3 | 6870 | 2.84 | 62 | Male | First manifestation of RA | MS |
| 21 | Rheumatoid Arthritis | Actemra | positive | positive | NS | NS | NS | NS | Female |  | MS |

|  | Disease | Treatment | RF | Anti-CCP | CRP | KL score | Age ( years) | Sex | Note | Used in |
| --- | --- | --- | --- | --- | --- | --- | --- | --- | --- | --- |
| 22 | Rheumatoid Arthritis | Abatacept, MTX | positive | positive | NS | NS | 60 | Female |  | WB |
| 23 | Rheumatoid Arthritis | Leflunomide | positive | positive | NS | NS | 68 | Female |  | WB |
| 23 | Rheumatoid Arthritis | Rinvoq, Leflunomid | negative | negative | NS | NS | 52 | Male |  | MS,WB |
| 24 | Rheumatoid Arthritis | Adalimumab, Leflunomid | negative | negative | NS | NS | 51 | Male |  | MS,WB |
| 25 | Rheumatoid Arthritis | NS | negative | positive | <0.3 | NS | 50 | Male |  | WB |
| 26 | Inflammatory Arthritis | No treatment | negative | negative | NS | NS | 67 | Male | = Patient 19 SynFluid | MS,WB |
| 27 | Osteoarthritis | NS | NS | NS | NS | 4 | 60 | Female |  | MS |
| 28 | Osteoarthritis | NS | NS | NS | NS | 4 | 57 | Female |  | MS |
| 29 | Osteoarthritis | NS | NS | NS | NS | 4 | 55 | Femle |  | MS, WB |
| 31 | Osteoarthritis | NS | NS | NS | NS | NS | 70 | Female | = Patient18 SynFluid | MS,WB |
| 32 | Osteoarthritis | NS | NS | NS | NS | 4 | 77 | Male |  | WB |

### Isolation of EVs via ultracentrifugation (UC)

Patient-derived synovial fluid was centrifuged 2000 x g for 10 min at RT aliquoted and stored at -80°C. A 1 ml aliquot was thawed on ice, pre-treated by addition of hyaluronidase (1 μg/ml Hyaluronidase-Type IV-S - Sigma) and protease/phosphatase inhibitors (Halt protease, Thermo Fisher Scientific) incubated for 45 min at 37 °C, subsequently diluted 1:5 in filtered PBS. Then, EVs were pelleted via ultracentrifugation (1h, 4 °C, 100.000 x g) using Optima XPN-90 ultracentrifuge (Beckman Coulter). Resulting pellets were taken up in 50 μl protease/phosphatase inhibitor supplemented RIPA buffer.

### Isolation of EVs via size exclusion chromatography (SEC) + Ultrafiltration

<u>A.  Synovial fibroblast- and THP1-derived EVs (from cell culture supernatant)</u>: For isolation of synovial fibroblast-derived EVs, 2.5 – 4 x 10^6^ synovial fibroblasts were cultured in a T175 flask for further growing and incubated for 72 h (37 °C, 5 % CO_2_). Afterwards, the medium was changed to exosome-depleted medium (Gibco) and the cells were incubated for 48 h releasing extracellular vesicles into the supernatant.

For the isolation of THP1-derived EVs, 30 x 10^6^ cells were primed with PMA (10 ng/ml) in a T175 flask and incubated overnight (37 °C, 5 % CO2). Then, the primed THP1 cells were stimulated for 24 h with LPS (100 ng/ml) and INFγ (20 ng/ml).

The implementation of the EV isolation for both cell types (THP1, synovial fibroblasts) was performed as followed:

Dead cells and cell debris were removed from the supernatant (30 ml) by centrifugation at 500 x g, 10 min and subsequently 3000 x g, 10 min. Afterwards, the supernatant was 15-fold (2ml end volume) concentrated and ultrafiltrated with Amicon Ultra centrifugal units (100 kDa cut off, Millipore).

The ultrafiltrated supernatant was transferred to a size exclusion column (qEV1, Gen2, 35nm, IZON) discarding the default buffer volume (4 ml = fractions 1-4), while collecting the EV containing fractions 5-9 (700 μl/fraction). Fractions 5-9 were pooled and concentrated/ultrafiltrated to a 14-fold lower volume using Amicon Ultra centrifugal units (10 KDa cut off, Millipore).

<u>B. Synovial fluid-derived EVs:</u> Patient-derived synovial fluid was centrifuged 2000 x g for 10 min at RT aliquoted and stored at -80°C. A 2 ml aliquot of synovial fluid was thawed on ice and pre-treated by addition of hyaluronidase (1 μg/ml Hyaluronidase-Type IV-S - Sigma), DNAse I (20 U/ml, Roche, 04536282001) and protease/phosphatase inhibitors (Halt protease, Thermo Fisher Scientific) incubated for 45 min at 37°C. The supernatant of a following centrifugation step (1000 x g, 5 min, RT) was filtered with a filter syringe (0.45 μM).

Before implementation of EV isolation via SEC (size exclusion column (qEV1, Gen2, 35nm, IZON), the pre-treated synovial fluid was diluted 1:2 with filtered PBS (0.22μm).

During SEC the default buffer volume (4ml = fractions 1-4) was discarded, while the EV containing fractions 5-9 (700 μl/fraction) were collected. Fractions 5-9 were pooled and concentrated to a 14-fold lower volume using Amicon Ultra centrifugal units (10 KDa cut off, Millipore).

The isolated EVs from both a) and b) were used for downstream analysis like Western Blot and for the establishment of the separation method using Miltenyi separation beads as described below. The particle concentration of the isolated EVs were analyzed with NTA (see below).

### Nanotracking Analyzer (NTA) for measurement of EV concentrations

The particle concentrations (particle/ml) of all SEC-isolated EVs were measured using a ZetaView PMX 110 V3 (ParticleMetrix). EV samples were diluted and added into the laser chamber, where the particle concentration is measured by analyzing the Brownian motion of each particle. NTA-software ZetaView (version 8.05.14 SP7) was used for evaluation of the data and determination of the particle concentration.

### Nano-FCM

The Nano-Analyzer (from Nanofcm) allows to measure the size (diameter) and concentration (EVs/ml) of EVs, but also the detection of stained surface marker on EVs. For specific surface marker detection, fluorophore-labelled antibodies (CD63_PE (BD Pharminogen-556020), CD90_PE (Miltenyi-REAffinity 130-114-903), Podoplanin_PE (Miltenyi-REAffinity 130-117-687), Isotype_PE (Miltenyi - REA Control,130-113-438)) were centrifuged 10 min, 17.000 x g at 4°C for removal of aggregates. Subsequently, 1 μl of undiluted antibody was mixed with 9 μl EVs (2x10^10^ EVs/ml) and incubated for 30 min at RT in the dark. For measurements samples were further diluted 1:100 (CD63) or 1:25 (CD90, PDPN). NP40 control was performed by adding 40 μl NP40 (2%) after staining incubation (1 μl antibody + 9 μl EVs). The NP40 approach was incubated again 30 min at RT in the dark and were then further diluted with filtered PBS to the final measurement concentration for NanoFCM.

### Negative stain transmission electron microscopy (TEM) of SEC-isolated EVs before and after bead separations

For TEM, 50 μl of SEC-isolated EVs in a concentration of 1x10^9^ EVs/ml and 50 μl of Eluate and Flowthrough (concentration unknown) of the CD90/PDPN bead separation were used. Grids (400-mesh copper grids with Parlodion carrier film (50 nm thick) and vapour-deposited carbon (10 nm thick)) were glow discharged before 5 μl of sample was added and incubated for 60 seconds. The grid is blotted with filter paper and subsequently 3 drops of ddH2O (50 μl each) were pipetted onto a parafilm for washing the grids on it (blotted with filter paper in between). For negative staining, two drops of 2% uranyl acetate aqueous solution were pipetted on parafilm, placing the grids on top (staining duration was 10 seconds), then again blotted with filter paper. Finally, grids were examined via TEM (CM100 Phillips).

### Dividing EVs and cell lysate proteins into cytosolic and membrane proteins (MemPER)

For both, synovial fibroblasts and their SEC-isolated EVs, we used the MemPER Plus Protein Exctraction KIT from Thermo Fisher Scientific (89842) according to the guidelines for dividing their protein content into cytosolic and membrane located proteins.

In short: 2.5 x 10^6^ synovial fibroblasts were harvested by scraping the cells in cold PBS, centrifugating them (300 x g, 10 min), then adding 375 μl permeabilization buffer supplemented with protease/phosphatase inhibitor.

2x10^10^ SEC-isolated EVs were precipitated overnight with an Exosome isolation KIT (Thermo Fisher Scientific – Total Exosome Isolation Reagent (from cell culture media)- 4478359) according to the manufacturers protocol. The resulting pellet was taken up in 200 μl permeabilization buffer supplemented with protease/phosphatase inhibitor. Both, EV and cell lysate protein was incubated 10 min at 4°C constantly shaking and subsequently centrifugated 16000 x g, 15 min. The supernatant was transferred into a new tube (= cytosolic fraction), while the pellets were resuspended in 250 μl (cell lysate) or 100 μl (EV) of solubilization buffer supplemented with protease/phosphatase inhibitor. Both were incubated 30 min at 4°C constantly shaking and again subsequently centrifugated 16000 x g, 15 min. The supernatant (=solubilized membrane and membrane-associated fraction) was collected into a new tube.

### Western Blot

SEC-isolated EVs and bead separated EVs were precipitated overnight with an Exosome isolation KIT (Thermo Fisher Scientific – Total Exosome Isolation Reagent (from cell culture media)-4478359) according to the manufacturers protocol. The resulting pellet was resuspended and lysed in RIPA-Buffer containing phosphatase and protease inhibitors. The protein amount was measured with Micro-BCA-Protein Assay KIT (Thermo Fisher Scientific-23235). 50 mM DTT and 1x Lämmli (final concentration) was added to all samples and were heated for 5 minutes at 70°C, before these were loaded on a 4-15 % SDS gradient gel (BioRad) for dividing the proteins according to their size. If not stated otherwise, we loaded all protein for eluates, washes and flowthroughs we received from our separation experiments.

The protein transfer was performed using TransBlot (BioRad). The membrane was blocked in blocking buffer (5 % milk solved in 0.05 % TBS-T) for 1 h, followed by three washing steps à 5 min in washing buffer (1xTBS-T - 0.05 % Tween20) and an overnight incubation of primary antibodies at 4 °C (CD90 (CST-13801S-Rabbit-1:1000), Podoplanin (CST-9047-Rabbit-1:1000), CD68 (UltraMAB-UM870047-Mouse-1:2000), CD63 (Invitrogen-10628D-Mouse-1:1000), CD9 (Invitrogen-10626D-Mouse-1:1000), CD81 (Invitrogen-10630D-Mouse-1:1000), CD163 (CST- 93498-Rabbit-1:1000), Syntenin (Abcam-ab133267-Rabbit-1:1000), Flotillin-1 (BD-610820-Mouse-1:1000), Annexin-1 (BD-610067-Mouse-1:5000), Calnexin (CST-2679S-Rabbit-1:1000), CD48 (CST-29499T-Rabbit-1:1000)). After three washing steps, secondary antibodies were solved in blocking buffer and incubated for 1 h at room temperature constantly shaking (Goat-anti-Rabbit (CST-6211234-1:3000), Horse-anti Mouse (CST-7076S-1:3000)). Antibody-stained membranes were washed three times and finally rinsed with 1xTBS. The signals resulting from addition of the chemiluminescence solution (Thermo Fisher Scientific - Pierce ECL Western Blotting Substrate - 32106) were detected with the Fusion Fx imager (Vilber).

Western Blot quantifications were performed from repeated Western Blots (n=3) and signal intensities (pixel density) for each protein signal were analysed using Fiji/ImageJ (Version 2.14.0/1.54f). The data was normalized to the corresponding measurement of Syntenin-1.

### Immunocapture of EVs for separation of subpopulations with MicroBeads

For all immunocapture experiments, we used magnetic bead coupled antibodies (CD90 prototype MicroBeads, Podoplanin prototype MicroBeads, Isotype prototype MicroBeads), which were kindly newly produced and provided by Miltenyi (Miltenyi Biotec, Bergisch-Gladbach, Germany). For additional control experiments we used the Pan-separation KIT from Miltenyi (Mix: CD63, CD9, CD81, EV Isolation KIT Pan, human - 130-110-912).

In general, bead separations were performed as described in Miltenyi guidelines. In brief, 50 μl of the bead-coupled antibodies (CD90 prototype beads or Podoplanin prototype beads or Isotype beads) was added to isolated EVs or synovial fluid (dependent on experiment). The EV/antibody mixture was incubated for 1 h at room temperature mildly shaking and subsequently put on the magnetic column allowing to collect flowthrough, wash and eluates 1 and 2 after each other (for eluate 2, the elution step was repeated, to check whether there is still remaining material in the column after the actual elution step).

Dependent on the experiments, this protocol included some addition’s: For bead separations in synovial fluid (spike in, separation from synovial fluid-derived EVs and direct separation from synovial fluid) we put the flowthrough a second time over a fresh magnetic column and pooled both eluates to reduce the number of remaining beads in the flowthrough and higher enrich the marker positive EVs in the eluate. Additionally, the resulting flowthrough and wash were put over SEC and subsequently ultrafiltrated/concentrated (Amicon 10K, Millipore, final volume approx. 500 μl).

Finally, the flowthrough, the wash and both eluates were precipitated using a precipitation KIT (Thermo Fisher Scientific – Total Exosome Isolation Reagent (from cell culture media)- 4478359) according to the companies’ recommendations. Beads were removed through sequential centrifugation steps (2x 30 min, 1x 10min, 10.000 x g). Protein amounts were determined by Micro-BCA-Protein Assay KIT (Thermo Fisher Scientific – 23235).

### A. CD90 prototype, PDPN prototype and isotype prototype beads in cell culture-derived EVs

1 – 2x 10^10^ of SEC-isolated and up-concentrated EVs were used for the immunocapture based separations. For the mixed populations (synovial fibroblast-derived EVs mixed to THP1-derived EVs) the EV concentration was doubled through the 1:1 mixture of EVs (final concentration: 2 – 4x 10^10^). The whole separation process using magnetic columns was performed as recommended in the Miltenyi guidelines.

Eluates, wash and flowthrough were precipitated (Thermo Fisher Scientific – Total Exosome Isolation Reagent (from cell culture media)- 4478359) and the resulting pellets were resuspended in 20 μl RIPA-buffer supplemented with phosphatase and protease inhibitors.

### B. Spiking human synovial fluid with synovial fibroblast-derived EVs from cell culture (CD90 prototype and Isotype prototype beads)

A thawed synovial fluid aliquot (500 μl) was pre-treated as described before (Isolation of EVs via SEC). For this experiment, we used a synovial fluid sample, where we couldn’t detect CD90^+^ EVs before on Western Blot. 500 μl pre-treated synovial fluid was spiked with 1 – 2x 10^10^ of SEC-isolated and up-concentrated synovial fibroblast-derived EVs from cell culture (500 μl) and filled up with filtered PBS to a final volume of 2 ml (viscosity-decrease of synovial fluid), which is then used for bead separation. The pellets after precipitation (Thermo Fisher Scientific – Total Exosome Isolation Reagent (from cell culture media)- 4478359) for flowthrough, eluates and wash were taken up in 30 μl protease/phosphatase inhibitor added RIPA.

### C. CD90 prototype and Podoplanin prototype beads applied in synovial fluid-derived material

Patient-derived synovial fluid was pre-treated as described before (Isolation of EVs via SEC). For that, 2ml pre-treated synovial fluid was transferred to SEC (qEV1, Gen2, 35nm, IZON) to isolate synovial fluid-derived EVs using the same procedure as described above. The resulting EVs (1ml volume, approx. 5x 10^10^ particles/ml) were used for the Podoplanin and CD90 prototype bead separation.

For direct use of beads in synovial fluid, 2 ml pre-treated synovial fluid was divided into 4 approaches, which were pooled again after the whole separation process: 500 μl pre-treated synovial fluid was diluted with 1500 μl filtered PBS (4x), before bead separation.

After bead separation and precipitation of EVs, the pellets of the resulting 4 eluates, 4 washes and 4 flowthroughs were pooled in 40 μl protease/phosphatase inhibitor added RIPA, respectively.

### Proteomics

#### CD90 bead separation in synovial fluid-derived EVs

CD90 bead immunocapture was performed from synovial fluid of arthritis patients (SEC pre-isolated). The lysis of EVs were performed direct on the magnetic column after washing steps. The columns were temporary closed during their location at the magnet and 100 μl lysis buffer (5% SDS, 50 mM Tris, pH=7.8) were added on top of the column. After an incubation time of 40 min (RT), columns were opened and the lysis buffer dropped through the column. Further 100 μl lysis buffer was added to the column. The flowthrough was precipitated (Thermo Fisher Scientific – Total Exosome Isolation Reagent (from cell culture media)- 4478359) and the pellets were resuspended in 200 μl lysis buffer. For removal of beads, flowthrough was centrifuged 2x 40 min, 10.000 x g at RT, transferring the supernatant into fresh Eppis. Both, eluate and flowthrough were heated 95°C for 5 min and stored at -20 °C until further preparation for Proteomics.

Eluates and Flowthrough were adjusted with SDS and TEAB (Triethylammonium bicarbonate) to the final concentration of 5% and 100 mM, respectively. Samples were reduced by addition of TCEP (Tris-2(carboxyethyl)phospine) to the final concentration of 10 mM and 10 min incubation at 95°C. Proteins were then alkylated with 20 mM iodoacetamide for 30 min at RT (protected from the light). Samples were then digested using S-Trap™ micro spin columns (Protifi) according to the manufacturer’s instructions. Shortly, 12 % phosphoric acid was added to each sample (final concentration of phosphoric acid 1.2%) followed by the addition of S-trap buffer (90% methanol, 100 mM TEAB pH 7.1) at a ratio of 6:1. Samples were mixed by vortexing and loaded onto S-trap columns by centrifugation at 4000 g for 1 min followed by three washes with S-trap buffer. Digestion buffer (50 mM TEAB pH 8.0) containing sequencing-grade modified trypsin (Promega) was added to the S-trap column and samples were incubated for 1h at 47 °C. Peptides were eluted by the consecutive addition and collection by centrifugation at 4000 g for 1 min of 40 μl digestion buffer, 40 μl of 0.2% formic acid and finally 35 μl 50% acetonitrile, 0.2% formic acid. Samples were dried under vacuum and stored at -20 °C until further use.

Dried peptides were resuspended in 0.1% aqueous formic acid, loaded onto Evotip Pure tips (Evosep Biosystems) and subjected to LC–MS/MS analysis using a Exploris 480 Mass Spectrometer (Thermo Fisher Scientific) fitted with an Evosep One (EV 1000, Evosep Biosystems). Peptides were resolved using a Performance Column – 30 SPD (150μm × 15cm, 1.5 um, EV1137, Evosep Biosystems) kept at 40 °C fitted with a stainless-steel emitter (30 μm, EV1086, Evosep Biosystems) using the 30 SPD method. Buffer A was 0.1% formic acid in water and buffer B was acetonitrile, 0.1% formic acid.

The mass spectrometer was operated in DIA mode. MS1 scans were acquired in centroid mode at a resolution of 120,000 FWHM (at 200 m/z), a scan ranges from 350 to 1500 m/z, AGC target set to Standard and maximum ion injection time mode set to Auto. MS2 scans were acquired in centroid mode at a resolution of 15,000 FWHM (at 200 m/z), precursor mass range of 400 to 900 m/z, quadrupole isolation window of 12 m/z without window overlap, a defined first mass of 120 m/z, normalized AGC target set to 3000 % and maximum injection time mode set to Auto. Peptides were fragmented by HCD (Higher-energy collisional dissociation) with collision energy set to 28% and one microscan was acquired for each spectrum.

The acquired raw-files were searched using the Spectronaut (Biognosys v19.0) directDIA workflow against a Homo sapiens database (consisting of 20,360 protein sequences downloaded from Uniprot on 2022/02/22) and 392 commonly observed contaminants. Default settings were used.

### Bioinformatics and Statistics

#### Data download and processing of publicly available data sets

We downloaded data sets provided by HPA, Uniprot, PanglaoDB and ExoCarta from the corresponding online repository/web resources. Published data sets were retrieved either from supplementary data provided by the publisher on the corresponding Journal/publication location or from official depositories. We compared all downloaded data sets on the basis of the HPA annotation score of 20’082 proteins (Status OCT2024). Therefore, we first matched all retrieved protein and gene names listed in the downloaded data sets to their corresponding gene/protein symbols annotated in HPA. Genes or proteins in the downloaded data sets not annotated in HPA were not included in the analysis.

Data were analyzed and bar plots/violin plots generated using GraphPad Prism (v.10.2.1) or Microsoft Excel. Data set overlaps were performed and Venn diagrams created using www.interactivenn.net^120^. To gain more insight into the molecular functions (MF), biological processes (BP), cellular composition (CC), and pathway enrichment of the identified SF-EV proteins, or the differentially expressed proteins in fraction E and FT, we performed GO enrichment and KEGG pathway analysis using g:Profiler ^121^.

### MS data processing, normalization strategy, PCA, differential expression analysis

#### Data processing

The raw Spectronaut output data was imported into R (R version 4.3.2 and Bioconductor version 3.18) and converted into a QFeature object for an intergrated analysis of feature to protein level measurements. This analysis included data filtering, aggreagtion and imputation steps^122^. First, the data was filtered to only include measurement calls with a protein group q-value < 0.01. Next, features were defined by the unique combination of peptide sequence, precursor charge, fragment ion and product charge. Each feature required at least 3 measurements across the entire data set. Features were aggregated to peptide level by summing all feature intensities for a given peptide per sample. Spectronaut reports two types of missed measurements which require different handling during data aggregation. Intensity values below 1 were treated as measurements below detection limit and were retained at the aggregated level if all feature intensities of a given peptide were below 1 in that sample. Unmeasured features (“NA” values) were retained at the peptide level if all feature intensties for a given peptide in that sample were missing. A next filtering step excluded sample-specific peptides, i.e. peptides which were only present in a single biological replicate of one sample group as defined by source (synovial fluid, synovial fibroblasts), fraction (flowthrough, eluate) and disease state. Peptide intensities were then log2-transformed with a pseudo-count of 1. Peptides with an intensity below detection limit (“0” values) were imputed by a Min-Prob strategy, which samples values from a Gaussian distribution with mean set on low intensity values in each sample. Imputation was done with the Qfeatures function impute and method=‘MinProb’ and otherwise default options. All “NA” peptide intensities were retained after imputation. In the next step, peptides were aggreagted to protein level by a median-polish strategy using the Qfeatures function aggregateFeatures and as method the MsCoreUtils function medianPolish^123^. The imputed protein levels still contained NA values for proteins unmeasured in a given sample. If protein measurements are missing across all samples of a given sample group they are most likely specifically absent in this condition. These cases were therefore treated by an additional MinProb-based imputation. All remaining NA values were retained in the down-stream differential analysis, as they can be handeled by limma^124^. In a last step, protein intensities per sample were median centered using the QFeatures function normalize and method ‘diff.median’. Principal component analysis (PCA) was performed with package pcaMethods using the method pca, which allowed the treatment of missing data with the nipalsPca algorithm^125^.

#### Differential expression analysis

The QFeature object was converted into an ExpressionSet object using the normalized protein level data. Cell surface protein scores for each protein were derived from the surfacegenie website (https://gundrylab.shinyapps.io/surfacegenie/)^77^. Differential expression analysis was baed on the limma package^124^. Briefly, a linear model was fit based on the four experimental groups built by crossing source (synovial fluid, synovial fibroblasts) and fraction (flowthrough, eluate). The patient effect was taken into account by estimating sample correlations within patient group samples using the duplicateCorrelation function of limma. Functions lmFit and eBayes were used to test specific contrasts such as eluate vs. flowthrough in synovial fluid samples. Significance was evaluated based on FDR-corrected P-values.

### Statistics

Descriptive statistics for continuous variables are presented as median and range, while proportions are expressed as percentages. Comparisons between samples were performed using Mann-Whitney U test with a Welch correction or student’s paired t-test. For multiple comparisons, statistical significance was determined using one-way analysis of variance (ANOVA) with Dunn’s correction, or two-way ANOVA with uncorrected Fishers’s LSD. P values < 0.05 were considered statistically significant. Data was processed in GraphPad Prism version 10.0 for MacOSX (GraphPad Software Inc). Where stated, outliers were identified and excluded using GraphPad Prism algorithm (representation of normalized proteomic data for exemplary proteins in Figure 7b).

## Acknowledgements

We would like to thank all enrolled patients, the medical teams and supporting personnel of the Department of Rheumatology of the University Hospital of Basel and the Blood Donation Center SRK beider Basel for sample provision, collection, and storage. In addition, we thank the Nano Imaging Lab of the Swiss Nanoscience Institute at the University of Basel, especially Susanne Erpel, for assessing TEM imaging. Calculations were performed at sciCORE (http://scicore.unibas.ch/) scientific computing center at University of Basel.

## Contributors

ANT and DK conceived the study, designed the experiments, wrote and approved the manuscript.

ANT performed experiments and data analysis required for this study

SK and ANT performed experiments and data analysis required for this study

SK wrote and approved the manuscript.

DK provided funding and interpreted clinical data.

EH, SHM and SG performed experiments, read and approved the manuscript.

KB performed MS measurement, read and approved the manuscript

FG and DB performed bioinformatic data analysis, and read and approved the manuscript

SW and UH provided prototype magnetic beads and material, read and approved the manuscript

## Patient and public involvement

Patients and/or the public were not involved in the design, or conduct, or reporting, or dissemination plans of this research.

## Ethics approval

The study was approved by the ethics committee of Northwest and Central Switzerland (EKNZ N° 2019-01693).

## Data availability statement

All data relevant to the study are available upon reasonable request and/or will be uploaded to an online data repository.

## Supplementary Figure legend

Supplementary Figure S1:

**Additional information regarding synovial fluid-derived samples.** (a): The total protein amount received from UC of 1ml synovial fluid varies prominently. Thus 50 μg of loaded protein is representing different percentages of total protein across the samples. (b) Percentage of surface marker distribution on EVs between different arthritis patient samples by assessing the specific EV marker signal intensity normalized to corresponding Syntenin-1 intensity in Western Blot.

Supplementary Figure S2:

**Nano Tracking Analyzer (NTA) and NanoFCM assessments of RA vs. OA synovial fibroblast-derived EVs** (a) NTA measurement curve exemplary for one measurement and additionally NTA measurement detecting size and concentration of synovial fibroblast-derived EVs (RA vs. OA) (b) NanoFCM assessment of synovial fibroblast-derived EVs (RA, OA) using CD63_PE, CD90_PE and PDPN_PE single staining - additional information data sheet. (c) Controls including CD63_PE only (without EVs), CD90_PE only (without EVs), PDPN_PE only (without EVs), NP40 CTRL, Isotype_PE staining, EVs only (without AB staining), and PBS only.

Supplementary Figure S3:

**Division of different western blot signal heights into two different Podoplanin populations and PAN bead separation.** (a) Shown are the Western Blot results for Podoplanin signal after Podoplanin bead separation of synovial fibroblast-derived EVs between 4 different patient samples. (b) PAN bead separation of synovial fibroblast-derived EVs. PAN beads (CD63, CD81, CD9). Eluate (E), Flowthrough (FT), Wash (W)

Supplementary Figure S4:

**EV isolation via SEC-UF coupled and direct immunocapture approach.** (a) TEM images of PDPN MicroBead separated synovial fluid-derived EVs (Eluate vs. Flowthrough). Compared are MicroBead separation between SEC pre-isolated EVs and the direct approach. White arrows point on EVs. (b) CD90 MicroBead separation in SEC pre-isolated, synovial fluid-derived EVs. A synovial fluid sample low expressing CD90 on EVs was used for MicroBead separation (CD90 negative for Western Blot detection of ultracentrifugated synovial fluid, CD90-negative for Western Blot detection of “bulk” SEC pre-isolated EVs). A weak CD90 signal could be detected in the Eluate after CD90 MicroBead separation via Western Blot. (c) MicroBead separation in concentrated cell culture supernatant without pre-isolation via SEC. (a) Graphical illustration and Western Blot results for EV isolation and MicroBead separation. CD90 and Podoplanin (PDPN) MicroBead separation direct from concentrated cell culture supernatant. Eluate (E), Flowthrough (FT), Wash (W), Cell lysate synovial fibroblasts (CL).

Supplementary Figure S5:

**Data comparisons with reference data sets, Principal component analysis (PCA) and differential expression analysis of synovial fluid EV proteins.** (a) Overlap of our data set with other reference data sets: First overlap with ExoCarta, a data set containing protein/RNA/lipid cargoes in small EVs. Second, a data set from Human protein atlas including a set of 551 high-abundant plasma proteins. (b) Pie charts showing the number of proteins in SF EV proteome significantly enriched in E (orange) vs FT (light blue) fraction assessed by differential expression analysis with adjusted p-value <0.05 and a Log2FC = +/-1.5. Number in grey area represent significantly enriched proteins but outside of the set Log2FC cut-off. (c) PCA of MS-measured Eluate (E) and Flowthrough (FT) protein samples derived from the prototype CD90-bead based immunocapture of SEC isolated EVs either from synovial fluid of arthritis patients (N= 11; E= green dots; FT= violet dots), or from supernatant of cultured synovial fibroblasts isolated from arthritis patients (N= 10; E= red dots; FT= blue dots). Circles in the corresponding colors visualize the clustering of the sample groups.

Supplementary Figure S6:

**Additional analysis generated by using the Broad Institute Single Cell Portal (SCP) with AMP Phase 1 project data set 1 Rheumatoid Arthritis.** (a) Venn diagram displaying all significantly enriched proteins in E *vs* FT (assessed by differential expression analysis), which are overlapping with the scRNA-Seq data sets representing synovial fibroblasts (Korsunsky *et al.*, 2022^79^) or arthritic synovial tissue cell populations (Edalat *et al.*, 2024^80^). (b) Gene set overlapping and gene expression projection assessments represented as Venn diagrams and dot blots using enriched proteins in E *vs* FT with SPC scores 1-0. Protein set was projected on CSP using AMP Phase I project RA Dataset (Zhang, F. *et al.*, 2019^39^). These figures display the complete results corresponding to Figure 9d, which showed the enriched proteins in E *vs* FT with SPC scores 4-3-2. (c) Spiderweb chart of Tabula Sapiens/CELLxGENE gene expression data for selected cell types compatible with the 11 CTS from the current study. Values represent the average fraction of cell type expressing the selected gene signature (=significantly enriched proteins in E *vs* FT with SPC score 4).

