## Supplementary figures and images for "Identification and proteomic profiling of CD90^+^ small EVs using a refined immunocapture separation approach targeting stromal-derived EV subpopulations in synovial fluid of arthritis patients"

### Supplementary Figure S1

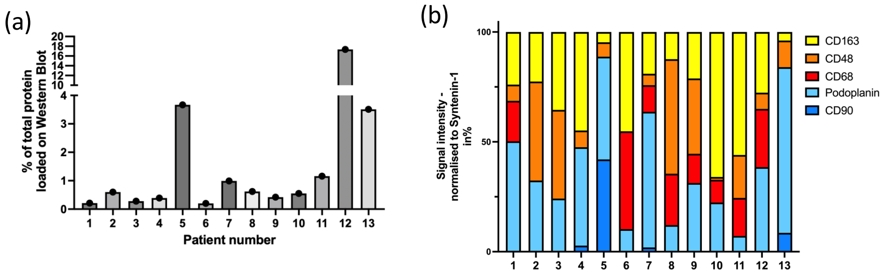

### Supplementary Figure S2

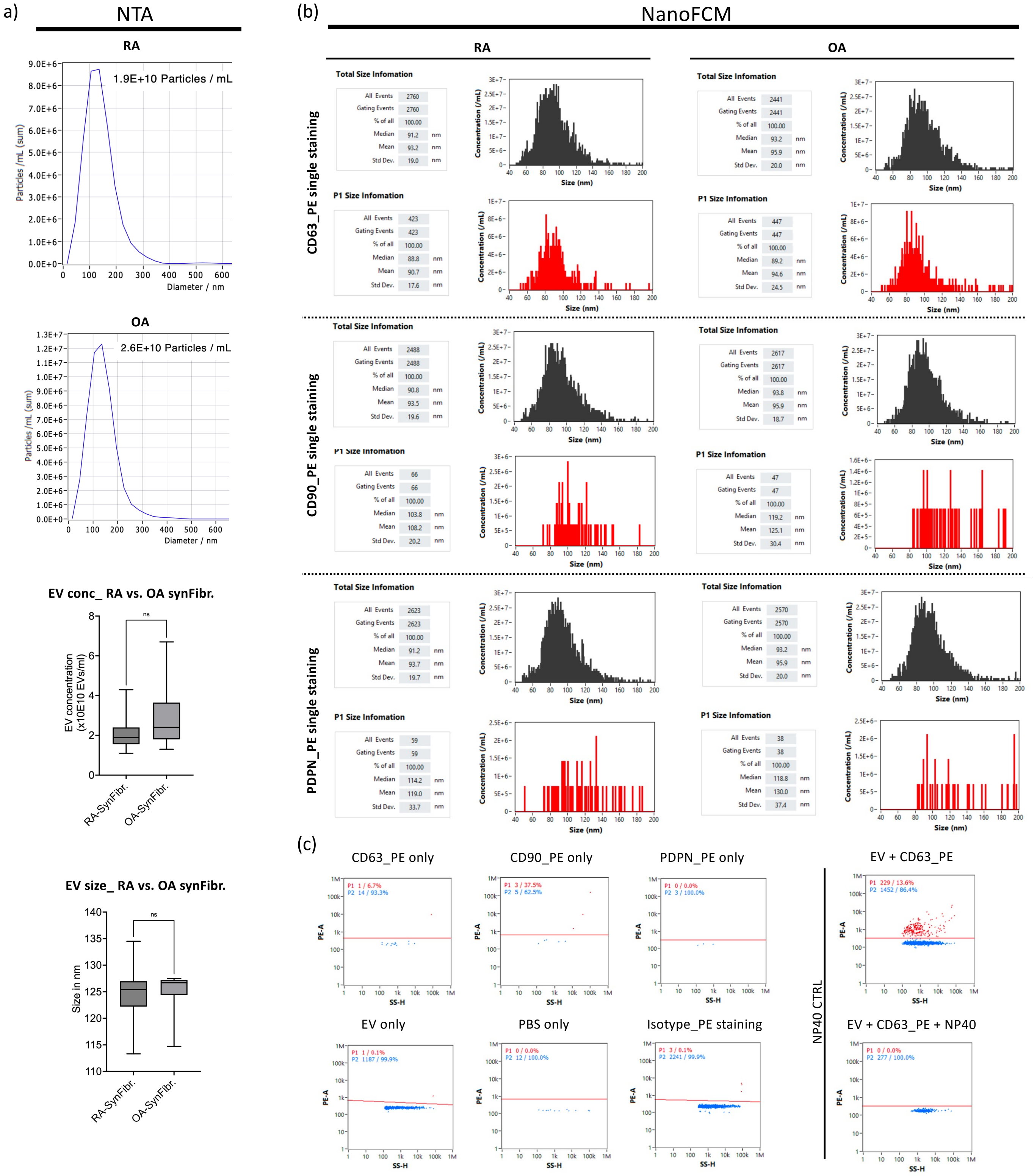

### Supplementary Figure S3

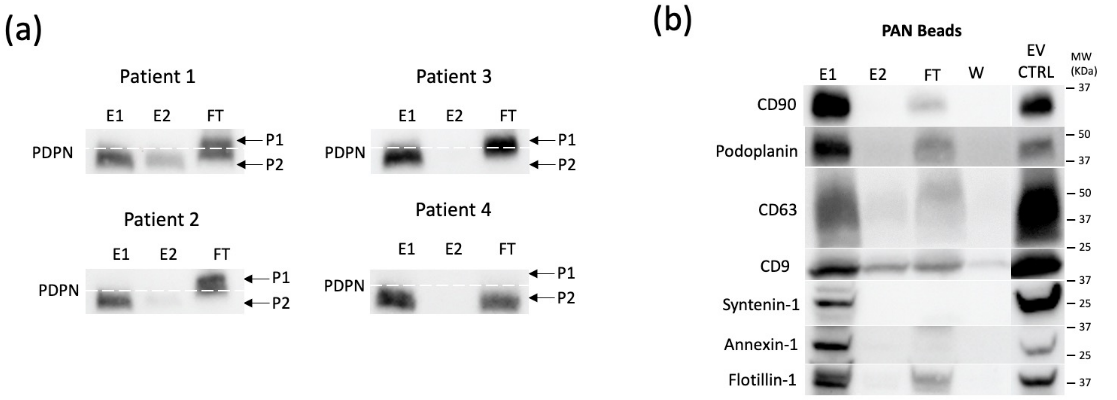

### Supplementary Figure S4

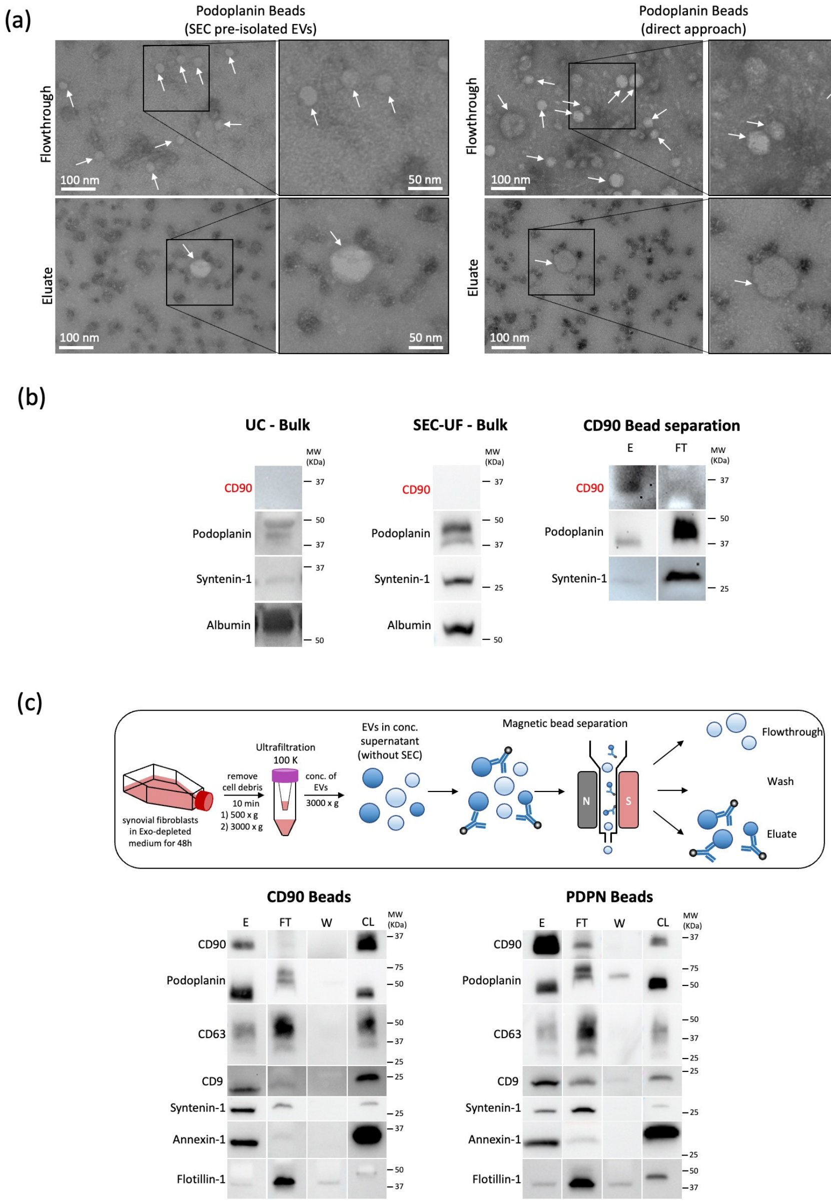

### Supplementary Figure S6

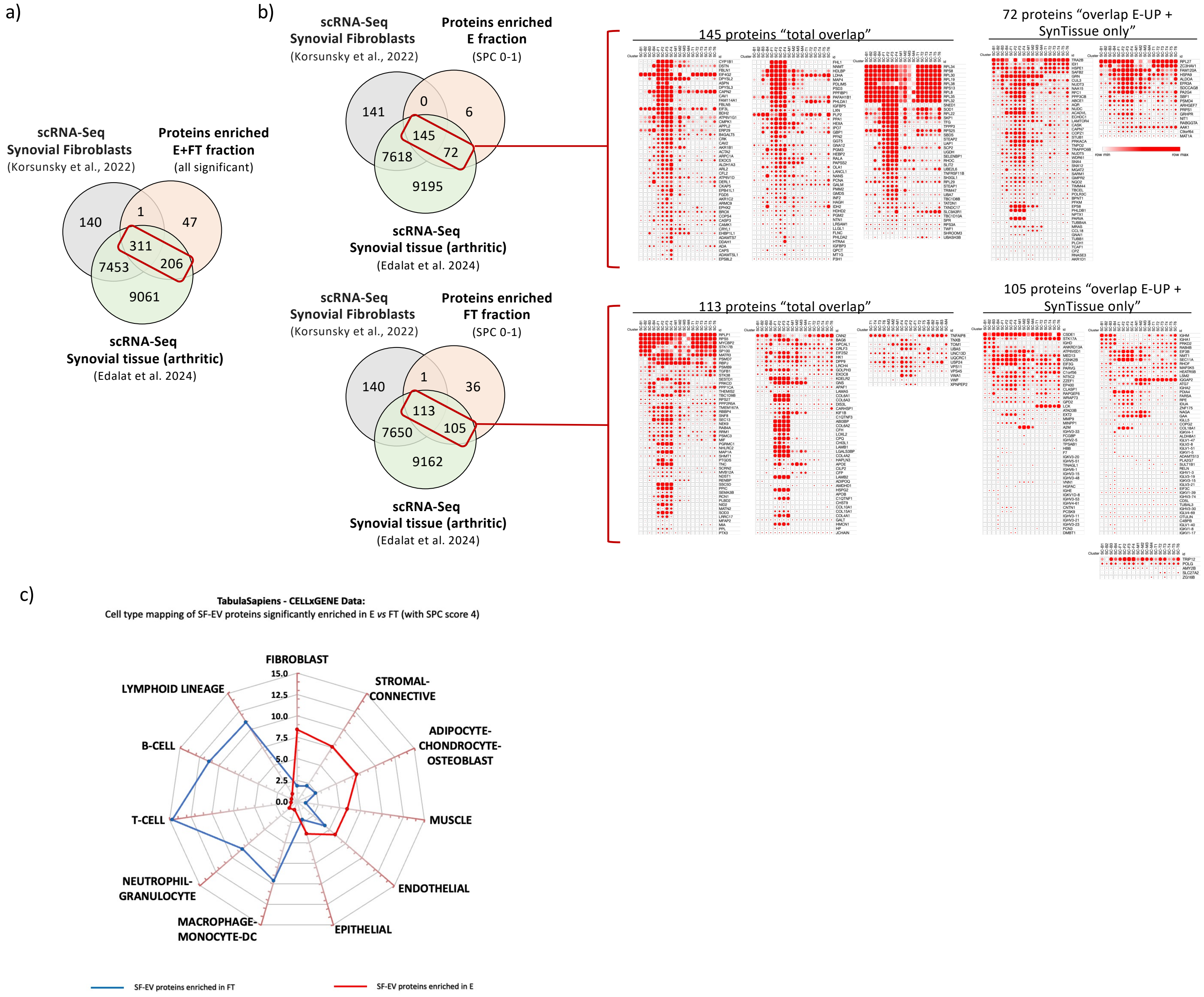
