## Supplementary Figure S5 for "Identification and proteomic profiling of CD90^+^ small EVs using a refined immunocapture separation approach targeting stromal-derived EV subpopulations in synovial fluid of arthritis patients"

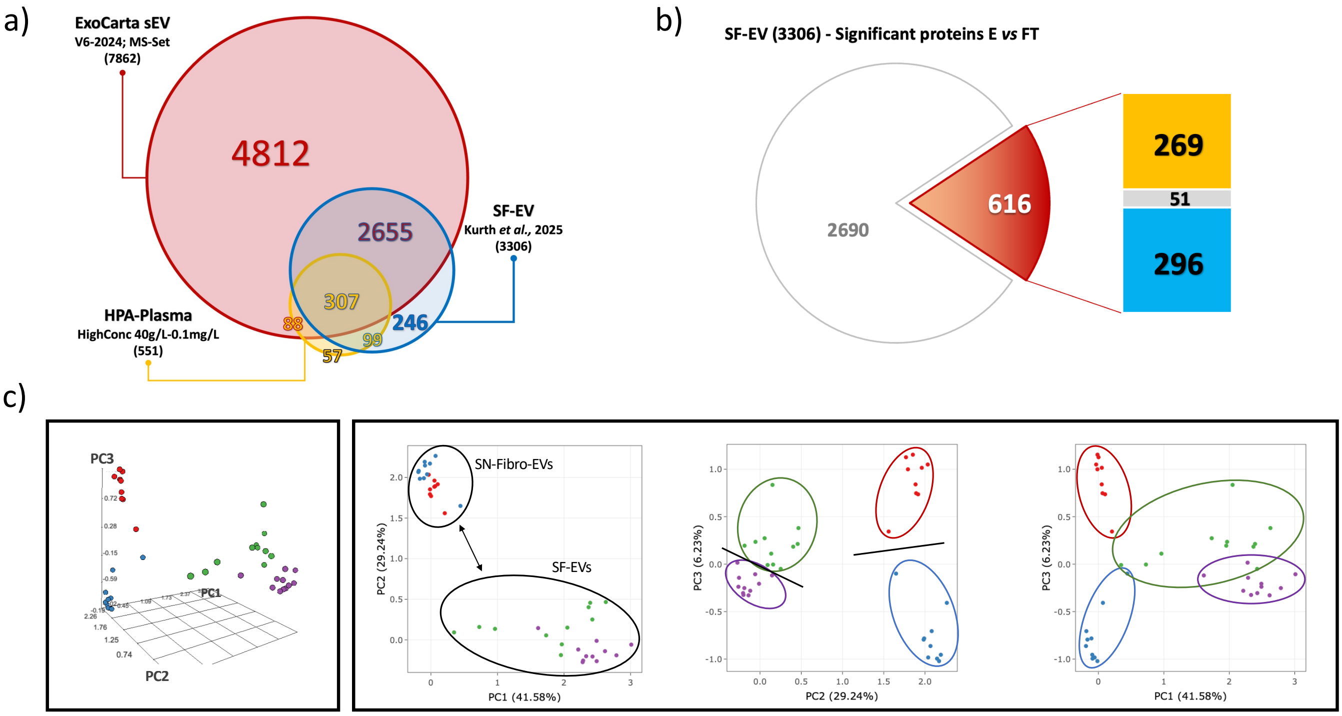


**Synovial fluid EV samples (SF-EVs; N=11)**

*Eluate fraction (separated CD90⁺EVs)*

*Flowthrough fraction*

**Cultured synovial fibroblast EV samples (SN-Fibro-EVs; N=10)**

*Eluate fraction (separated CD90⁺EVs)*

*Flowthrough fraction*
